# Coincident epithelial signals restrain commensal-specific CD8αβ^+^ T cells in the intestine

**DOI:** 10.64898/2026.04.29.720676

**Authors:** Tayla M. Olsen, Matthew J. Dufort, Sheenam Verma, Venkata Krishna Kanth Makani, Benjamin Cameron, Addison R. Gralen, Aaron J. Johnson, Alok V. Joglekar, Allyson L. Byrd, Adam Lacy-Hulbert, Oliver J. Harrison

## Abstract

The intestinal epithelium harbors a large population of microbiota-dependent CD8αβ^+^ T cells whose antigen specificity and regulation are ill-defined. By identifying MHCIa-restricted TCRs and generating tetramers against the gut commensal Segmented Filamentous Bacteria, we demonstrate that a single commensal species drives a clonally expanded, antigen-specific CD8αβ^+^ T cell within the intraepithelial lymphocyte compartment. Mechanistically, the intestinal epithelium coordinates coincident signals governing this population: peptide:MHC-dependent TCR engagement drives pIEL accumulation, while αvβ6-mediated TGFβ activation restraints effector cell differentiation. Perturbation of epithelial cell-mediated TGFβ activation diverts commensal-specific CD8^+^ T cells toward inflammatory differentiation states transcriptionally convergent with those observed in ulcerative colitis. The intestinal epithelium thus functions as a dual-signal organizer of commensal-specific CD8^+^ T cell responses, coupling differentiation to restraint through spatially coincident molecular cues.

## Main Text

The microbiota is essential for development and education of the immune system and tissue homeostasis, yet poses a persistent challenge to the host: commensal-specific immune recognition must be tightly restrained to prevent inflammation at barrier surfaces.^1–3^ In the intestine, diverse bacterial taxa, including Segmented Filamentous Bacteria (SFB), *Clostridia*, *Helicobacter*, *Bacteroidetes*, and *Akkermansia muciniphila* among others elicit robust antigen-specific CD4^+^ T cell responses that shape intestinal immunity,^1–14^ demonstrating that commensals actively instruct adaptive immune differentiation programs. Beyond lamina propria CD4^+^ T cells, the intestinal epithelium contains a specialized compartment of T cell receptor (TCR)αβ^+^ peripherally induced intraepithelial lymphocytes (pIEL) positioned in intimate contact with intestinal epithelial cells (IECs). CD8αβ^+^ pIEL are nearly absent in germ-free mice and accumulate markedly upon colonization,^4,15,16^ yet unlike their CD4^+^ counterparts, far less is known about which defined commensal species elicit cognate CD8αβ^+^ pIEL responses, and the cellular and molecular mechanisms governing such responses remain undefined.

A central challenge of host-microbiota mutualism is the need to balance antigen-specific immune recognition with durable tolerance.^17^ Commensal-specific T cells must be restrained under homeostatic conditions to prevent inappropriate tissue damage, while retaining the capacity to respond during infection or tissue injury.^1,18–20^ When such restraint fails, pathological immune activation ensues, a hallmark of chronic inflammatory disorders including inflammatory bowel disease (IBD), in which inflammatory pIEL have been observed.^21–24^

Tissue-resident lymphocytes are shaped by local cues reflecting the unique structural, metabolic, and microbial features of each barrier site.^25–29^ In the intestinal epithelium, CD8αβ^+^ pIEL are exposed to epithelial-derived signals distinct from those in lymphoid organs or the lamina propria, which critically determine their differentiation, function, and tolerance.^14,21,30,31^ Among these, the MHCIb molecule Thymic Leukemia (TL) antigen, highly expressed on IECs, raises the threshold for TCR activation through its interaction with CD8αα.^32,33^ How epithelial signals are integrated to maintain antigen-experienced CD8^+^ T cells in a non-pathogenic state, particularly in the context of commensal colonization and continuous microbial antigen exposure, remains an open question. Locally derived immunoregulatory signals within the epithelial microenvironment must play a central role in this process, yet their identity and cellular source are poorly understood.

TGFβ is a central immunoregulatory cytokine essential for immune tolerance and limiting immunopathology, particularly in the intestine;^34–37^ and a compelling candidate for such a role. Unlike freely diffusible cytokines, it is produced as a latent complex requiring spatially controlled activation. Which cell types locally activate TGFβ within the intestinal microenvironment, and how this signal is directed toward CD8αβ^+^ pIEL remains to be determined. Here, by developing commensal-specific MHCIa-restricted TCRs and tetramers we identify SFB as a driver of antigen-specific CD8αβ^+^ pIEL responses in the small intestine and show that cognate peptide MHC (pMHC) interactions with IECs are required for their differentiation and continuous TCR stimulation within the epithelium. Despite harboring potent effector potential, SFB-specific CD8αβ^+^ pIEL are actively restrained, a state whose epithelial and cell-intrinsic basis we define here.

## Results

### SFB colonization induces accumulation of clonally expanded CD8ɑβ^+^ pIEL

To determine how the commensal microbiota influences the CD8αβ^+^ pIEL compartment, we compared C57BL/6 mice harboring distinct microbial communities. Wild-type (WT) mice from Taconic Biosciences (Tac) exhibited increased frequencies and total numbers of CD8αβ^+^ pIEL relative to their counterparts from Jackson Laboratories (Jax) **(Fig. 1A)**. This expansion arose post-weaning, was vertically transmissible, and unlike the transient accumulation of lamina propria (LP) CD8αβ^+^ T cells following weaning, was stable and persisted into adulthood **(Fig. S1A-B)**.

**Fig. 1.**
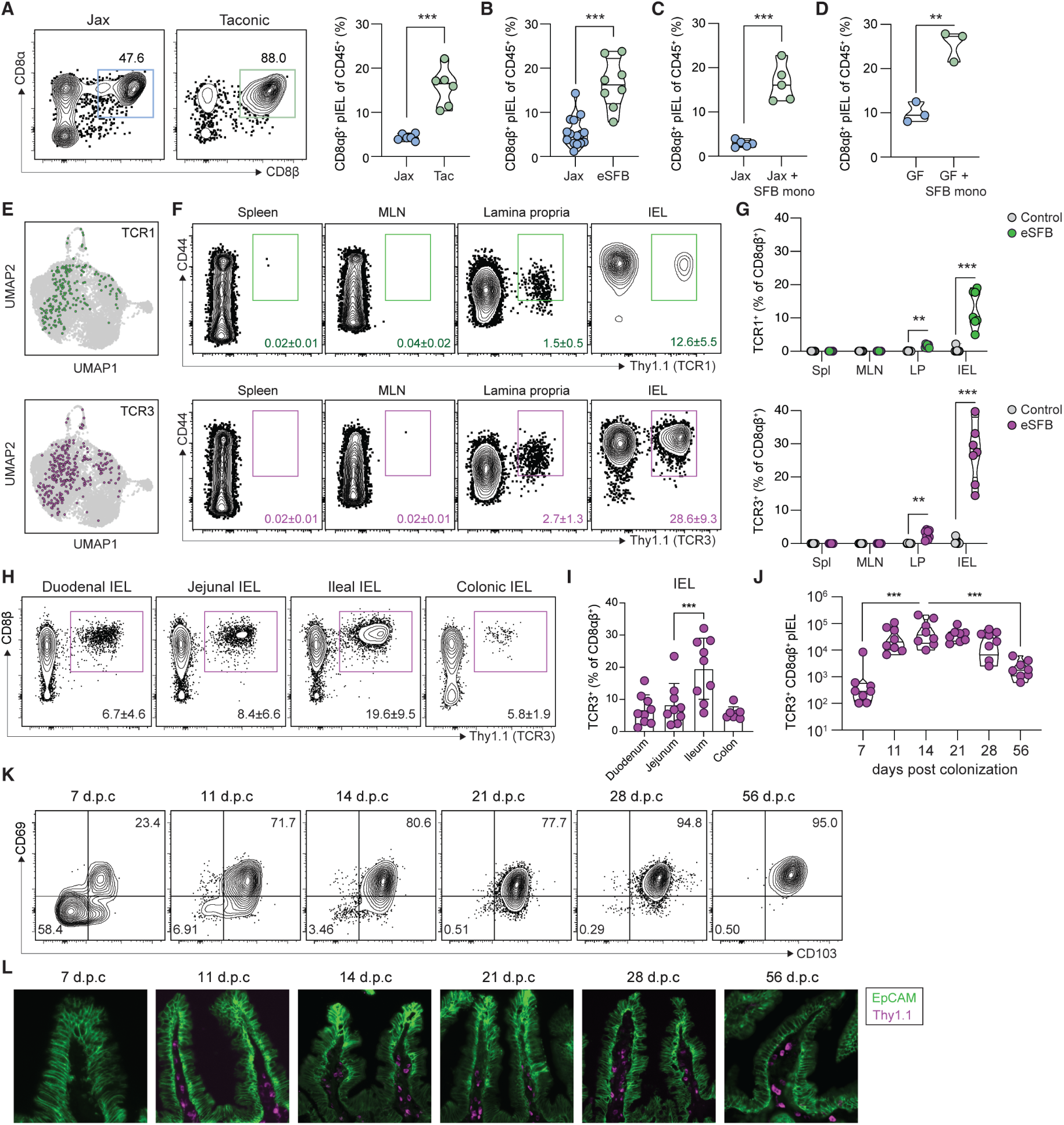
SFB colonization induces accumulation of clonally expanded CD8ɑβ^+^ pIEL. **(A)** Representative flow plots of CD8αβ^+^ pIEL pre-gated on live CD45^+^Thy1.2^+^TCRβ^+^CD4^-^ and violin plot of CD8αβ^+^ T cells as a percent of total live CD45^+^ cells from the IEL fraction of the last 10cm of the small intestine of mice from Jax and Tac, n=6 pooled from two independent experiments. **(B)** Violin plot of CD8αβ^+^ T cells as a percent of total live CD45^+^ cells from Jax mice and Jax mice colonized with eSFB 28 days post colonization (d.p.c.), n=8-15 per group pooled from three independent experiments. **(C)** Violin plot of CD8αβ^+^ T cells as a percent of total live CD45^+^ cells from Jax mice and Jax mice colonized with SFB from monoassociated feces 28 d.p.c., n=5 per group. **(D)** Violin plot of CD8αβ^+^ T cells as a percent of total live CD45^+^ cells from germ-free B6 mice colonized with SFB from monoassociated feces 28 d.p.c., n=3 per group. **(E)** Uniform manifold approximation and projection (UMAP) plot highlighting the cluster position and frequency of TCR clonotypes TCR1 and TCR3 from 10x single cell V(D)J sequencing of CD8αβ^+^ pIEL from two Jax mice and two H2-M3^-/-^ mice 28 d.p.c. with eSFB. **(F)** Representative flow plots from eSFB colonized Jax mice and **(G)** violin plot of retrogenic cells as a frequency of total CD8αβ^+^ T cells from Jax mice receiving 10,000 naive TCR1^Rg^ (green) and TCR3^Rg^ CD8^+^ T cells (pink) with or without eSFB analyzed 14 d.p.c and T cell transfer, n=7-8 per group from two independent experiments. **(H)** Representative flow plots and **(I)** Bar graph of TCR3^Rg^ CD8^+^ T cells as a frequency of total CD8αβ^+^ T cells from Jax mice receiving 10,000 naive cells with eSFB analyzed 14 d.p.c. and T cell transfer. The small intestine was segmented in thirds, and the colon was taken along with the cecum, n=9 mice per group from two independent experiments. **(J)** Violin plot of total TCR3^Rg^ CD8^+^ T cells and **(K)** their flow plots showing CD69 by CD103 from Jax mice receiving 10,000 naive cells with eSFB analyzed over time, n=8 per group from two independent experiments. **(L)** Small intestine imaging from Swiss Rolls of TCR3^Rg^ CD8^+^ T cells from Jax mice receiving 10,000 naive cells with eSFB analyzed over time, stained with anti-Thy1.1 (magenta) and Ep-CAM (green). P values were calculated using (A-D) unpaired t test (G-J) ordinary one-way ANOVA with Tukey’s multiple comparisons, and (K) quadrant frequencies plus/minus the standard deviation are shown. Statistical significance denoted as ** P <0.01, *** P <0.001, and **** P <0.0001.

Segmented Filamentous Bacteria (SFB) colonizes mice from Tac but not Jax^38^ **(Fig. S1C)** and elicits robust CD4^+^ T cell and B cell responses in the gut.^38–42^ We therefore tested whether SFB colonization alone induced CD8αβ^+^ pIEL accumulation using fecal transfer from enriched SFB (eSFB) donors.^39,40^ De novo colonization with eSFB flora promoted CD8αβ^+^ pIEL accumulation in the small intestinal ileum **(Fig. 1B, S1D)**, as did colonization with SFB monoassociated feces of SPF Jax and germ-free C57BL/6 or Swiss Webster mice **(Fig. 1C-D, S1E)**, demonstrating that a single commensal is sufficient to drive establishment of intestinal epithelial-resident CD8^+^ T cells in genetically distinct mouse strains. Consistent with these cells being SFB-specific, mesenteric lymph node (MLN)-derived CD8ɑβ^+^ T cells from eSFB-colonized mice produced IFN-ɣ in response to SFB antigens ex vivo **(Fig. S1F)**, corroborating recent reports.^4^ To directly assess antigen specificity of SFB-induced CD8αβ^+^ pIEL, we conducted single-cell RNA (scRNAseq) and V(D)J sequencing. Given that microbe-specific MHCIb-restricted CD8^+^ T cells contribute to tissue repair and intestinal immunity,^18,43,44^ we assessed CD8αβ^+^ pIEL from WT and *H2-M3*^-/-^ mice.^45^ Robust clonal expansion in both genotypes indicated that H2-M3-restricted cells are not major contributors to the SFB-induced CD8αβ^+^ pIEL pool **(Table S1)**. We then retrovirally expressed the ten most clonally expanded TCRαβ heterodimers in WT CD8^+^ T cells and adoptively transferred them into congenic recipients subsequently colonized with control or eSFB flora **(Fig. 1E, S1G, Table S1)**. Two TCRs, TCR1 and TCR3, drove robust CD8αβ^+^ pIEL accumulation in SFB colonized hosts, which we prioritized for further study **(Fig. S1H-I)**.

To enable further investigation of TCR1 and TCR3 and their ability to differentiate into CD8αβ^+^ pIEL, we generated TCR retrogenic (TCR^Rg^) mice using a modified “On Time” system that produces naive CD8αβ^+^ T cells expressing a TCR of interest and a truncated Thy1.1 marker.^46,47^ Upon adoptive transfer, naive TCR1^Rg^ and TCR3^Rg^ CD8αβ^+^ T cells became activated in hosts de novo colonized with eSFB, and accumulated in the spleen, MLN, LP and intestinal epithelium, with the latter harboring the highest frequency of TCR3^Rg^ CD8αβ^+^ T cells **(Fig. 1F-G)**. TCR3^Rg^ CD8αβ^+^ pIEL, but not LP cells, were further enriched within the ileum **(Fig. 1H-I, S1K)**, consistent with the restricted anatomic niche of SFB at this site.^38^ Located primarily in small intestinal villi, TCR3^Rg^ cells progressively acquired a CD69^+^CD103^+^ phenotype from 7 days post-colonization (d.p.c.) and remained a significant fraction of the pIEL compartment for up to 2 months, detectable by both flow cytometry and imaging **(Fig. 1J-K)**. CD69^+^CD103^+^ differentiation was accompanied by upregulation of CD8αα, detectable by TL-tetramer staining **(Fig. S1L)**, a hallmark of pIEL differentiation.^11,21,30,33^ Together, these data establish that a single commensal species can drive clonal expansion and epithelial residency of antigen-specific CD8αβ^+^ T cells, a previously uncharacterized arm of the adaptive immune response to the intestinal microbiota.

### SFB-specific CD8αβ^+^ pIEL are MHCIa-restricted and recognize a unique SFB epitope

To determine the antigen specificity of TCR1^Rg^ and TCR3^Rg^ cells, we co-cultured them with purified antigen-presenting cells (APCs) and SFB-derived antigens. Both TCRs were activated by FACS-purified SFB, but not germ-free or Jax feces **(Fig. 2A, S2B)**,^38^ confirming that responses were driven by SFB-derived peptide antigens. Furthermore, TCR activation was dependent upon MHCIa-mediated antigen presentation, as *H2-Kb^-/-^H2-Db*^-/-^ APCs failed to activate either TCR **(Fig. 2A, S2B)**. Using DC2.4 cell lines deficient for individual MHCIa molecules,^48^ we determined that TCR1 is H2-Db-restricted, whereas TCR3 is H2-Kb-restricted **(Fig. 2B, S2A, S2C, S2D)**, demonstrating that both MHCIa molecules in C57BL/6 mice can present SFB antigens to drive commensal-specific CD8αβ^+^ pIEL differentiation.

**Fig. 2.**
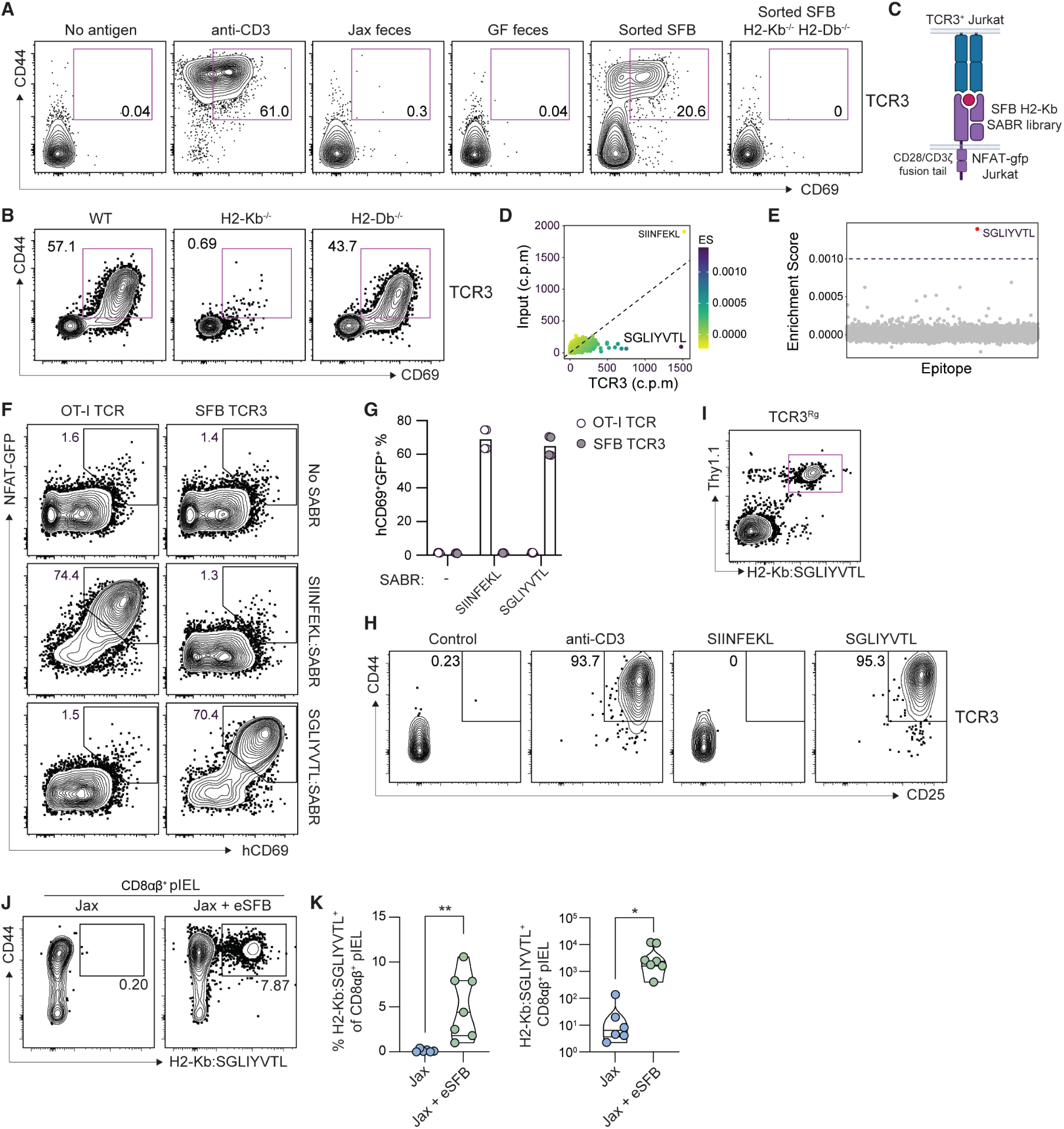
SFB-specific CD8αβ^+^ pIEL are MHCIa-restricted and recognize a unique SFB epitope. **(A)** Representative flow plots from co-cultured naive TCR3^Rg^ CD8^+^ T cells and purified CD11c^+^ WT or MHCIa^-/-^ antigen-presenting cells (APCs) treated with either anti-CD3, feces from Jax or germ-free mice, or FACS-purified SFB from monoassociated feces and incubated for 72 hours. Data are representative of two independent experiments. **(B)** Representative flow plots from co-cultured naive TCR3^Rg^ CD8^+^ T cells and DC2.4 APC cell lines WT or deficient for individual MHCIa molecules treated with eSFB feces and incubated for 72 hours. Data were confirmed with two independent experiments. **(C)** Schematic of Signaling and antigen-presenting bifunctional receptors (SABRs) epitope screening in Jurkat cells. **(D)** Scatter plot comparing normalized counts per million (cpm) epitope sequence reads from both SABR input library and sorted SABR cells cultured with Jurkat cells expressing TCR3. Calculated enrichment score is expressed as a heatmap statistic and **(E)** scatter plot of enrichment score versus each 15,769 epitopes from SFB-H2-Kb-SABR library sorted alphabetically. (D) and (E) were calculated from two independent library transduction events with sorted cells from triplicate co-cultures; SGLIYVTL appeared in all library samples. **(F)** Representative flow plots for single SABR co-cultures of OTI -and TCR3-expressing Jurkat cells and SABR-SIINFEKL and SABR-SGLIYVTL Jurkat cells and **(G)** the bar graphs of 2 replicates from two independent experiments from (F). **(H)** Representative flow plots from co-cultured naive TCR3^Rg^ CD8^+^ T cells and purified CD11c^+^ WT APCs treated with either anti-CD3 or 1µg/mL of each peptide and incubated overnight. Data were confirmed with two independent experiments. **(I)** Flow cytometry plot of splenic CD8^+^ T cells from SFB-free Thy1.1^+^ TCR3 retrogenic mice stained with SGLIYVTL-tetramer and analyzed by flow cytometry. **(J)** Representative flow plots and **(K)** violin plots of total numbers of CD44^+^SGLIYVTL-tetramer^+^ CD8αβ^+^ T cells and as a frequency of total CD8αβ^+^ T cells from Jax mice receiving 10,000 naive cells with eSFB, analyzed 14 d.p.c. and T cell transfer, n=7 per group pooled from two independent experiments. P values were calculated using (G) ordinary one-way ANOVA with Tukey’s multiple comparisons and (K) unpaired t test. Statistical significance denoted as * P <0.05, ** P <0.01, and *** P <0.001.

Given the magnitude of TCR3 responses relative to TCR1, we focused on TCR3 for epitope identification. Using an H2-Kb signaling and antigen-presenting bifunctional receptor (SABR) platform **(Fig. 2C, S2E)**^49–51^, we screened a library of 15,769 in-silico-predicted H2-Kb-binding peptides derived from the SFB-Mouse-NYU proteome against TCR3-expressing Jurkat cells (Methods for full details). This approach identified a single high-confidence epitope, SGLIYVTL **(Fig. 2D-E)**, derived from SFBNYU_003240, a putatively secreted protein containing a domain of unknown function (DUF4214) implicated in cell surface-anchoring.^52,53^ TCR3 specificity for SGLIYVTL was confirmed using a single SGLIYVTL-H2-Kb-SABR construct and in vitro stimulation of naive TCR3^Rg^ CD8^+^ T cells with recombinant SGLIYVTL or control SIINFEKL peptides **(Fig. 2G, 2H)**. TCR3^Rg^ CD8^+^ T cells were also detectable by H2-Kb:SGLIYVTL tetramer staining **(Fig. 2I)** and, following SFB colonization in vivo, H2-Kb:SGLIYVTL tetramer positive cells accumulated in the intestinal epithelium of SFB colonized mice where they recapitulated the CD69^+^CD103^+^ phenotype of TCR3^Rg^ CD8αβ^+^ pIEL **(Fig. 2J-K, S2F)**. Together, these data establish SGLIYVTL as an H2-Kb-restricted SFB epitope, validate the H2-Kb:SGLIYVTL tetramers for tracking endogenous commensal-specific CD8^+^ T cells in vivo, and now incorporate MHCIa-restricted CD8^+^ T cell responses into the suite of adaptive immune responses elicited by the commensal microbiota.

### SFB-specific CD8αβ^+^ pIEL accumulation is dependent upon epithelial cell antigen presentation

SGLIYVTL-specific TCR3^Rg^ and polyclonal endogenous CD8^+^ T cells accumulated in the intestinal epithelium in an SFB-dependent manner **(Fig. 1F, 2J-K)**. SFB colonization promoted IEC expression of H2-Kb **(Fig. S3A)**,^54,55^ raising the possibility that local TCR stimulation by IECs drives microbiota-specific CD8αβ^+^ pIEL expansion or retention. To test this, we adoptively transferred naïve Nur77^GFP^ TCR3^Rg^ CD8^+^ T cells and colonized mice with SFB. At 10 days post colonization (d.p.c.), SFB-specific CD8^+^ T cells expressed Nur77^GFP^ equivalently across the MLN, LP and IEL, consistent with recent TCR stimulation of clonally expanding cells **(Fig. S3B, S3C)**. By contrast, 28 d.p.c., when CD8αβ^+^ pIEL accumulation had stabilized **(Fig. 1J)**, Nur77^GFP^ expression was selectively enriched in pIEL relative to MLN- and LP-derived cells **(Fig. 3A-C)**, demonstrating ongoing TCR stimulation within the epithelial compartment, even in CD8αα-expressing cells. Notably, this distinguishes MHCIa-restricted SFB-specific CD8αβ^+^ pIEL from MHCII-restricted CD4^+^CD8αα^+^ pIEL which progressively lose TCR-responsiveness upon epithelial entry,^11^ and pathogen-specific pIEL, in which TCR signaling after pathogen clearance is not observed.^37,56^ EdU incorporation confirmed ongoing proliferation of TCR3^Rg^ pIEL, consistent with receipt of sufficient TCR stimulation to drive cell cycle entry **(Fig. 3D)**.

**Fig. 3.**
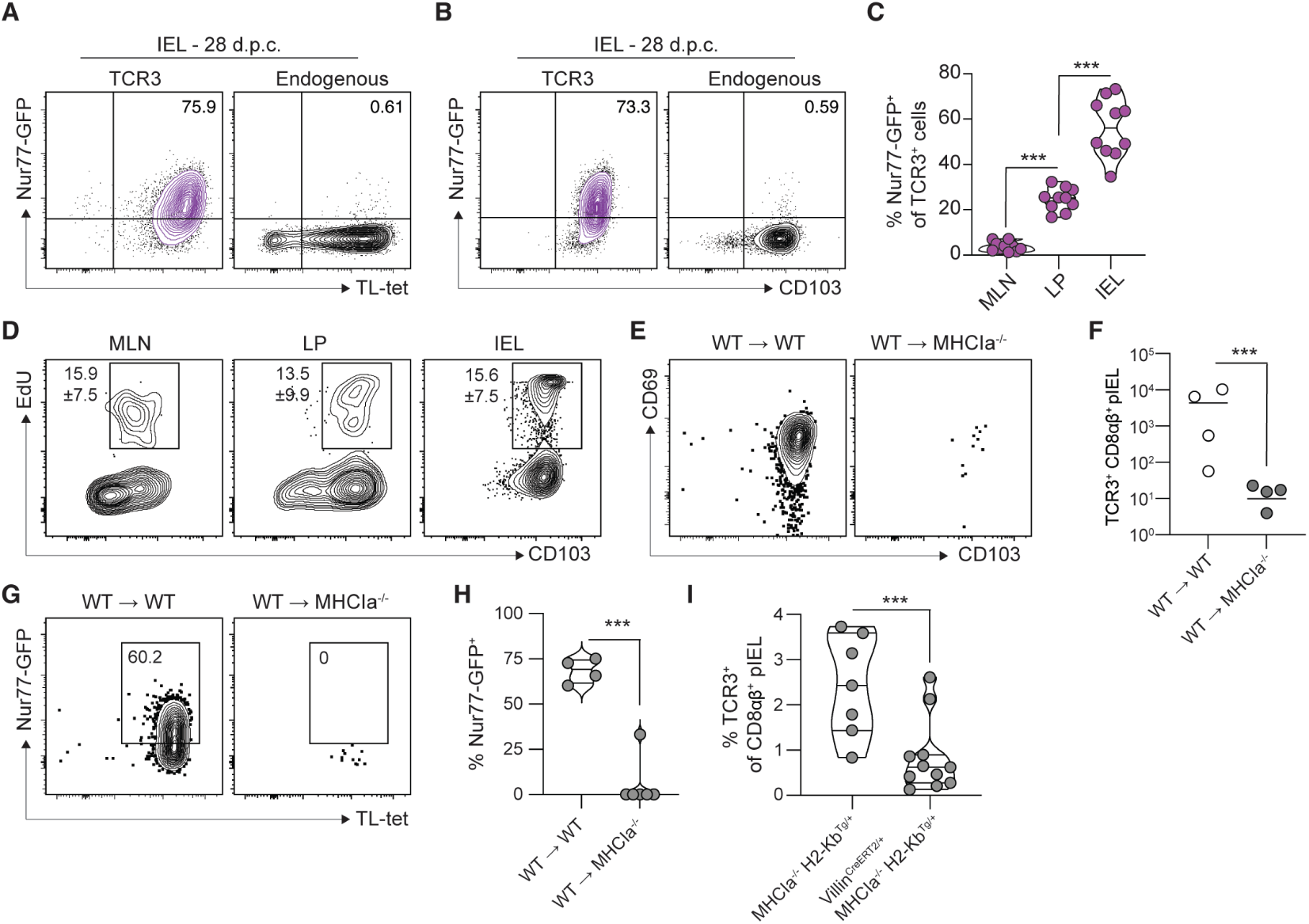
SFB-specific CD8αβ^+^ pIEL differentiation is dependent upon epithelial cell antigen presentation. **(A)** Representative flow plots showing TL-tetramer (CD8αα) staining and **(B)** CD103 staining of Nur77^GFP^ TCR3^Rg^ CD8^+^ or endogenous CD8αβ^+^ T cells from ileum of eSFB colonized Jax mice 28 d.p.c. and T cell transfer, n=8-10 per group pooled from two independent experiments. **(C)** Violin plot of Nur77^GFP^ as a frequency of TCR3^Rg^ CD8^+^ T cells from (A). **(D)** Representative flow plots of EdU incorporation in TCR3^Rg^ CD8^+^ T cells as frequency of total TCR3^Rg^ CD8^+^ T cells from mice 24 hours post EdU administration (0.5mg/mouse) analyzed 28 d.p.c. and T cell transfer, n=8 per group pooled from three independent experiments. **(E)** Representative flow plots showing CD69 and CD103 staining and **(F)** dot plot of total numbers of TCR3^Rg^ CD8^+^ pIEL, and TCR3^Rg^ CD8^+^ pIEL frequency of total T cells, and **(G)** representative flow plots showing Nur77^GFP^ by TL-tetramer (CD8αα), and **(H)** violin plot of Nur77^GFP+^ TCR3^Rg^ CD8^+^ T cells as a frequency of total TCR3^Rg^ CD8^+^ T cells present in WT or MHCIa^-/-^ bone marrow chimeras that received WT transplantation. Mice were analyzed 28 d.p.c. and T cell transfer (administered 2 months post chimerism), n=4-5 per group. **(I)** Violin plot of TCR3^Rg^ CD8^+^ pIEL as frequency of total CD8^+^ T cells in H2-Kb^Tg-fl/fl^ (MHCIa^-/-^) crossed to *Villin*^CreERT2/+^ administered with 4mg tamoxifen orally 4 and 1 day prior to T cell transfer and eSFB colonization, then 2, 5, and 8 d.p.c and T cell transfer. Mice were analyzed 14 d.p.c. and T cell transfer, n=7-11 per group pooled from two independent experiments. P values were calculated using (C and D) ordinary one-way ANOVA with Tukey’s multiple comparisons and (F, H and I) unpaired t test. Statistical significance denoted as not significant (ns), * P<0.05, ** P <0.01, *** P <0.001, and **** P <0.0001.

To determine whether IEC antigen presentation was required for these responses, we generated bone marrow chimeras in which WT hematopoietic cells were transplanted into WT or MHCIa^-/-^hosts. Loss of MHCIa on radio-resistant cells, including IECs, almost entirely ablated TCR3^Rg^ CD8αβ^+^ pIEL accumulation, and the few remaining pIEL lacked Nur77^GFP^ expression entirely **(Fig. 3E-H)**, demonstrating that CD8αβ^+^ pIEL accumulation and homeostatic TCR signaling both require IEC-derived MHCIa expression. To confirm this finding, we used inducible epithelial-specific H2-Kb deletion, by generating mice in which H2-Kb is the sole MHCIa molecule, that can be conditionally deleted from IECs by tamoxifen-mediated Villin^CreERT2^ activation (H2-Kb^Tg-flox^ x MHCIa^-/-^ x Villin^CreERT2/+^ mice).^57^ Epithelial H2-Kb-deletion both prior to and during eSFB colonization impaired TCR3^Rg^ CD8αβ^+^ pIEL accumulation relative to H2-Kb-competent controls **(Fig. 3I)**. Thus, IEC MHCIa antigen presentation is required for commensal-specific CD8αβ^+^ pIEL accumulation.

### Epithelial entry imprints a restrained phenotype on commensal-specific CD8αβ^+^ pIEL

At odds with blunted TCR responses in MHCII-restricted CD4^+^CD8ɑɑ^+^ pIEL, and those CD8ɑβ^+^ pIEL elicited by pathogen infection,^21,35,37,56,58–60^ SFB-specific CD8ɑβ^+^ pIEL demonstrate robust TCR-signaling under homeostasis **(Fig. 3A-D)**. Yet TCR3^Rg^ cells isolated from the IEL, unlike those from the LP, were more refractory to ex vivo TCR driven cytokine production with agonistic anti-CD3, despite retaining strong IFN-ɣ production capacity upon PMA/Ionomycin stimulation that bypasses proximal TCR signaling **(Fig. 4A)**. This dissociation between homeostatic TCR activity and ex vivo TCR responsiveness suggested that the epithelial microenvironment qualitatively remodels TCR signaling in commensal-specific pIEL. To identify the underlying signals contributing to this local immune regulation, we performed scRNAseq of TCR3^Rg^ CD8αβ^+^ T cells from the spleen, MLN, LP, and IEL of SFB-colonized mice. Louvain clustering resolved 10 transcriptionally distinct clusters **(Fig. 4B-C),** with spleen and MLN cells predominating in clusters 1-2 and IEL-enriched cells segregating into clusters 6, 7 and 8 **(Fig. 4D)**. Slingshot pseudo-time inference^61^ revealed five differentiation trajectories consistent with progressive tissue entry from secondary lymphoid organs through LP into IEL **(Fig. 4E)**. Splenic and MLN cells were enriched for lymphoid egress genes (*Sell*, *Klf2*, *S1pr1*), while LP and IEL cells upregulated gut homing and retention programs (*Itga4*, *Ccr9*, *Hic1*, *Cd69*) **(Fig. S4A-B)**, reflecting gut imprinting defined in other experimental systems.^29,62–64^ Within tissue-derived cells, pIEL expressed higher levels of *Gzmb* and *Itgae* than their LP counterparts, consistent with being pIEL markers in infectious settings **(Fig. S4C)**,^29^ along with canonical pIEL effector and residency genes (*Gzma, Gzmk, Ccl5*, *Itga1*, *Cd101*, *Cxcr6*)^29,65^ **(Fig. S4D)**.

**Fig. 4.**
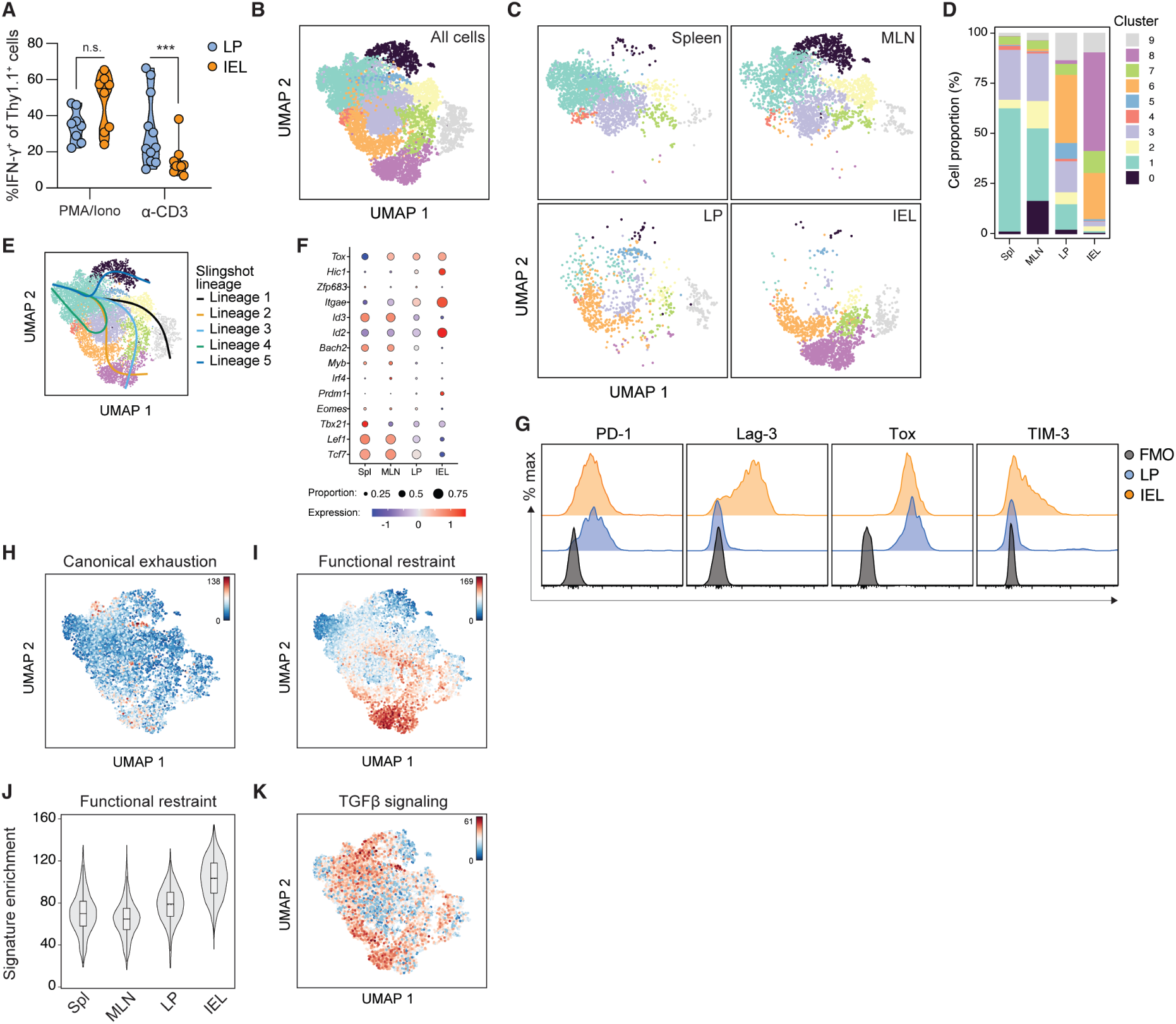
Epithelial entry imprints functional restraint on commensal-specific CD8αβ^+^ pIEL. **(A)** Violin plot of IFN-ɣ^+^ TCR3^Rg^ CD8^+^ T cells following *ex vivo* restimulation with PMA/ionomycin (2.5 hours) or anti-CD3 (5 hours) of cells from the ileum of Jax mice 14 d.p.c. and T cell transfer. **(B and C)** UMAP projection and **(D)** tissue Cluster distribution of 10x scRNAseq analysis of sorted TCR3^Rg^ CD8^+^ T cells from the spleen, MLN, ileum LP and IEL of Jax mice 11 d.p.c. and T cell transfer. Data were generated from cells pooled from 10 mice. **(E)** Slingshot cell lineage and pseudo-time inference analysis of differentiation trajectories of cells and **(F)** heatmap and proportion analysis of selected genes across tissues in cells from (B). **(G)** Protein expression of selected inhibitory receptors on TCR3^Rg^ CD8^+^ T cells from LP and IEL cells from Jax mice 14 d.p.c. and T cell transfer. **(H)** Canonical exhaustion and **(I)** functional restraint gene set enrichment projected on 10x scRNAseq data from (B) and **(J)** enrichment score violin plot for functional restraint gene set from projection in (I) plotted for spleen, MLN, LP and IEL. **(K)** TGFβ signaling gene set enrichment score projected on 10x scRNAseq data from (B). P values were calculated using (A) two-way ANOVA with Šídák’s multiple comparisons. Statistical significance denoted as not significant (ns), *** P <0.001.

SFB-specific CD8⍺β^+^ pIEL expressed transcription factors and coinhibitory molecules associated with terminally differentiated and/or exhausted T cells (*Tox, Prdm1*, *Lag3*, *Havcr2*, *Pdcd1*, *Tigit*, *Ctla4*, *Cd160*, *Cd244a*, *Cd39*, *Cd73*, *CD200R,* and *Cish*),^66–71^ of which only a subset were shared with LP-derived cells **(Fig. 4F-G, S4E-F),** pointing to compartment specific regulatory signals. Despite this profile, SFB-specific CD8⍺β^+^ pIEL were not enriched for a canonical T cell exhaustion signature **(Fig. 4H)**.^72^ Instead, they bore the transcriptional signature of “restrained” autoreactive islet-specific CD8^+^ T cells, that maintain functional potential despite coinhibitory receptor expression **(Fig. 4I-J)**.^72^ This restraint signature was largely confined to pIEL, implicating local epithelial signals in its induction. While Lag3 promoted functional restraint in autoreactive pancreatic CD8^+^ T cells,^72^ *Lag3* deletion did not impact CD8⍺β^+^ pIEL phenotype **(Fig. S4G)**.

Among candidate epithelial signals, pIEL were selectively enriched for a TGFβ-response gene signature **(Fig. 4K)**, and retained low-level expression of T stem-cell memory (Tscm) transcripts (*Tcf7*, *Slamf6*, *Lef1*) **(Fig. S4H-I)**, consistent with recent evidence that TGFβ and retinoic acid cooperate to protect pathogen-elicited pIEL from terminal effector differentiation.^73^ As such, in addition to myriad roles for TGFβ-signaling in establishment, maintenance, and local positioning of intestinal CD8^+^ T cells elicited by pathogen infection,^74–77^ these data point to epithelial TGFβ as a central enforcer of the restrained pIEL program and raise the question of how local TGFβ activation impacts commensal-specific CD8^+^ T cells within the epithelium.

### Epithelial cell ɑvβ6-dependent TGFβ activation restrains commensal-specific CD8ɑβ^+^ T cell effector function and reinforces pIEL differentiation

To determine the T cell-intrinsic requirement for TGFβ-receptor signaling in restraint of SFB-specific CD8ɑβ^+^ pIEL effector functions, we performed CRISPR-Cas9-mediated *Tgfbr2* deletion in TCR3^Rg^ CD8^+^ T cells prior to adoptive transfer and eSFB colonization. This resulted in accumulation of aberrant CD103⁻KLRG1^+^ CD8^+^ T cells in the MLN, LP, and IEL which was not observed in *Cd19* or *Itgae* targeting controls, consistent with reports that impaired TGFβ signaling during tissue entry drives terminal effector differentiation, and that CD103-E-cadherin ligation alone is not required for conventional pIEL differentiation **(Fig. 5A)**.^36,73,74,78^ Aberrant adaptive immune responses to commensal microbes are a hallmark of IBD.^23,79–81^ Notably, patients with ulcerative colitis (UC) harbor autoantibodies targeting epithelial integrin ɑvβ6, a key pathway for activation of latent TGFβ in vivo.^82–87^ To define how this epithelial axis impacts commensal-specific CD8ɑβ^+^ pIEL differentiation, we adoptively transferred TCR3^Rg^ CD8^+^ T cells into *Itgav*^fl/fl^ Villin^Cre/+^ mice, which lack αv-dependent integrins on IECs, prior to eSFB colonization. Epithelial *Itgav* deletion reduced CD103 expression in ∼50% of TCR3^Rg^ CD8⍺β^+^ pIEL, consistent with TGFβ-dependent *Itgae* regulation **(Fig. 5B)**.^36,88^ Transcriptional changes extended well beyond *Itgae*, however, as both CD103^+^ and CD103^-^ cells in *Itgav*^fl/fl^ *Villin*^Cre/+^ mice exhibited elevated expression of effector associated genes, including *Klrg1*, *Eomes*, *Ifng*, *Bhlhe40* and *Il12rb2*, relative to CD103^+^ pIEL from *Itgav*^fl/fl^ *Villin*^Cre/-^ controls **(Fig. S5A-B)**. Additionally, aberrant CD103^-^KLRG1^-^ and CD103^-^KLRG1^+^ TCR3^Rg^ CD8^+^ T cells accumulated to higher levels in MLN, LP and IEL of *Itgav*^fl/fl^ *Villin*^Cre/+^ mice **(Fig. 5C, S5C)**. Epithelial *Itgav* deletion also drove TCR3^Rg^ CD8^+^ T cell hyperplasia in the MLN and LP and resulted in alterations in terminal ileum villi architecture consistent with intestinal inflammation resembling that found in human IBD patient samples **(Fig. 5D, S5D).**^22^

**Fig. 5.**
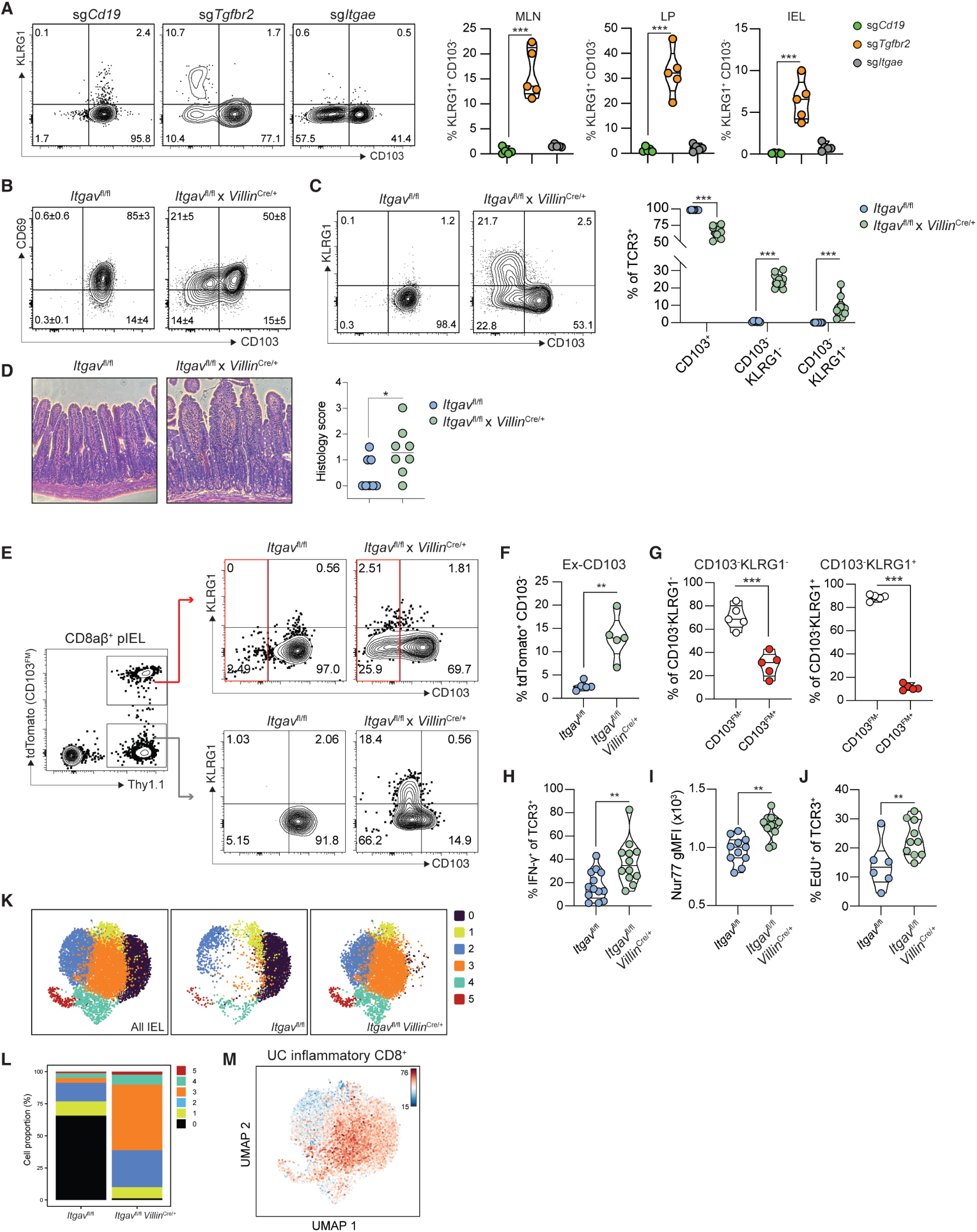
Epithelial cell ɑvβ6-dependent TGFβ activation restrains commensal-specific CD8ɑβ+ T cell effector function and reinforces pIEL differentiation. **(A)** Representative flow plots showing staining of KLRG1 by CD103 in TCR3^Rg^ pIEL knocked down with CRISPR-Cas9 RNPs of indicated gene and violin plots of CD103^-^KLRG1^+^ TCR3^Rg^ CD8^+^ T cells as a frequency of total TCR3^Rg^ cells in MLN, LP and IEL from Jax mice 14 d.p.c. and T cell transfer, n=10 per group pooled from two independent experiments. **(B)** Representative flow plots showing CD69 and CD103 staining on TCR3^Rg^ CD8^+^ pIEL from *Itgav*^fl/fl^ *Villin*^Cre/+^ or *Itgav*^fl/fl^ littermate control mice 28 d.p.c. and T cell transfer, n=7-10 per group pooled from two independent experiments. **(C)** Representative flow plots of KLRG1 and CD103 staining in TCR3^Rg^ pIEL and violin plot of CD103 and KLRG1populations of TCR3^Rg^ pIEL in mice from (B). **(D)** Representative images of hematoxylin and eosin staining of the last 1cm of the terminal ileum and dot plot of pathology scores in *Itgav*^fl/fl^ *Villin*^Cre/+^ or *Itgav*^fl/fl^ littermate control mice 28 d.p.c. and TCR3^Rg^ CD8^+^ T cell transfer, n=8 per group from two independent experiments. **(E)** Flow gating scheme and representative plots pre-gated CD44^+^CD8αβ^+^ T cells from CD103-CreERT2 fate mapping CD8^+^ T cells transduced *ex vivo* with TCR3 prior to transfer into *Itgav*^fl/fl^ *Villin*^Cre/+^ or *Itgav*^fl/fl^ littermate control mice. Mice were given 4mg tamoxifen every other day starting 12 d.p.c. and T cell transfer and analyzed day 21, n=5 per group. **(F)** Violin plot of CD103^-^tdTomato^+^ cells from *Itgav*^fl/fl^ *Villin*^Cre/+^ or *Itgav*^fl/fl^ littermate control mice from (E) as a frequency of total TCR3^Rg^ pIEL. **(G)** Violin plot of frequency of CD103^-^KLRG1^+^ TCR3^Rg^ CD8^+^ pIEL that are tdTomato^-^ or tdTomato^+^ from *Itgav*^fl/fl^ *Villin*^Cre/+^ mice from (E). **(H)** Violin plot of IFN-ɣ^+^ TCR3^Rg^ CD8^+^ pIEL following *ex vivo* restimulation with 1µg/mL anti-CD3 (5 hours) from the ileum of *Itgav*^fl/fl^ *Villin*^Cre/+^ or *Itgav*^fl/fl^ littermate control mice 14 d.p.c. and T cell transfer, n=12-13 per group pooled from three independent experiments. **(I)** Violin plot of the geometric mean fluorescence intensity (gMFI) of GFP from Nur77^GFP^ TCR3^Rg^ CD8^+^ pIEL of *Itgav*^fl/fl^ *Villin*^Cre/+^ or *Itgav*^fl/fl^ littermate control mice 28 d.p.c and T cells transfer, n=11-12 per group pooled from two independent experiments. **(J)** Violin plot of EdU^+^ TCR3^Rg^ CD8^+^ pIEL as a frequency of total TCR3^Rg^ CD8^+^ T cells from *Itgav*^fl/fl^ *Villin*^Cre/+^ or *Itgav*^fl/fl^ littermate control mice 28 d.p.c and T cells transfer, n=7-9 per group pooled from two independent experiments. **(K)** UMAP projection and **(L)** tissue Cluster distribution of 10x scRNAseq analysis of sorted TCR3^Rg^ CD8^+^ T cells from the pIEL of *tgav*^fl/fl^ *Villin*^Cre/+^ or *Itgav*^fl/fl^ littermate control mice 28 d.p.c. and T cell transfer. Data were generated from cells pooled from 3 mice for each genotype. **(M)** Functional restraint gene set enrichment projected on 10x scRNAseq data from (K). P values were calculated using (A) ordinary one-way ANOVA with Tukey’s multiple comparisons, (B) quadrant frequencies plus/minus the standard deviation are shown, (C, E, F, and G) two-way ANOVA with Šídák’s multiple comparisons, and (H-J) unpaired t test. Statistical significance denoted as * P<0.05, ** P <0.01, *** P <0.001.

Loss of epithelial *Itgav* gave rise to two transcriptionally distinct CD103⁻ populations, CD103⁻KLRG1^+^ and CD103⁻KLRG1⁻ cells, whose developmental origins were unclear. Since TGFβ signaling suppresses KLRG1 expression during effector differentiation,^89^ these populations could represent either cells that had previously acquired CD103 and subsequently downregulated it, or cells that had never initiated conventional pIEL differentiation. To distinguish between these possibilities, we used a fate-mapping approach to trace the origin of CD103^-^ pIEL, we adoptively transferring TCR3-transduced CD8^+^ T cells from CD103 fate-mapping mice (*Itgae*^CreERT2/+^ x *Rosa26*^LSL-tdTomato/+^)^56^ into *Itgav*^fl/fl^ or *Itgav*^fl/fl^ *Villin*^Cre/+^ mice. Ex-CD103^+^ (CD103^-^tdTomato^+^) cells were enriched in the LP and IEL but not MLN of *Itgav*^fl/fl^ *Villin*^Cre/+^ mice, accounting for ∼30% of CD103^-^KLRG1^-^ cells, whereas this population was not observed in control recipients **(Fig. 5E-F, S5E)**. By contrast, CD103^-^KLRG1^+^ cells were almost entirely tdTomato^-^ **(Fig. 5G, S5F)**, identifying them as a lineally distinct population that never acquired CD103, likely failing to initiate conventional pIEL differentiation in the absence of epithelial TGFβ. These cells accumulated not only in the IEL but also in the MLN and LP **(Fig. S5C)**, suggesting systemic dissemination of aberrantly differentiated commensal-specific CD8^+^ T cells. Together, these fate-mapping data reveal that impaired IEC αvβ6-dependent TGFβ activation subverts SFB-specific CD8^+^ T cells into divergent inflammatory lineages.

Loss of epithelial *Itgav* also unleashed effector function within SFB-specific CD8ɑβ^+^ pIEL: more TCR3^Rg^ CD8^+^ T cells from the IEL, but not LP, of *Itgav*^fl/fl^ *Villin*^Cre/+^ mice produced IFN-ɣ upon TCR restimulation, pIEL exhibited elevated Nur77^GFP^ expression in CD103^+^ cells and exhibited increased cellular proliferation, indicating heightened TCR responsiveness and loss of functional restraint **(Fig. 5H-J, S5G-H)**. ScRNAseq analysis of TCR3^Rg^ CD8ɑβ^+^ pIEL from *Itgav*^fl/fl^ *Villin*^Cre/+^ mice revealed a transcriptionally distinct cluster, Cluster 3, that was strongly enriched compared to *Itgav*^fl/fl^ controls **(Fig. 5K-L)**, and aligned transcriptionally with inflammatory CD8^+^ T cell subsets identified in human UC patients samples **(Fig. 5M)**,^22^ suggesting that loss of epithelial αvβ6-mediated TGFβ activation in mice recapitulates features of human intestinal inflammation associated with dysregulated CD8^+^ T cell responses. Together, these findings identify the intestinal epithelium as an active controller of antigen-specific CD8^+^ T cell responses to the microbiota, in which coincident pMHC presentation and αvβ6-mediated TGFβ activation restrain cells induced by commensal colonization, and whose dysregulation is associated with intestinal inflammation in both mice and humans.

## Discussion

Intraepithelial CD8αβ^+^ T cells expand in response to microbial colonization, yet whether specific commensals elicit cognate, antigen-driven responses in this compartment, and how such responses are regulated, has remained unknown. Here, using commensal-specific MHCIa-restricted TCRs and tetramers we reveal that SFB colonization drives a clonally expanded, antigen-specific CD8αβ^+^ T cell population of intraepithelial lymphocytes, establishing that a defined commensal species elicits a cognate CD8^+^ T cell response within the intestinal epithelium. These tools further reveal that the epithelium itself is a key regulator of this response: IEC pMHC presentation is required for pIEL accumulation and continuous TCR engagement, while coincident αvβ6-activated TGFβ enforces compartment-specific functional restraint. That both signals originate from the same cell type ensures that differentiation and restraint are spatially and mechanistically coupled, a logic that, when disrupted, results in inflammatory effector programs in commensal-specific CD8^+^ T cells transcriptionally aligned with those found in patients with IBD.

A key finding is that commensal-specific CD8ɑβ^+^ pIEL actively signal through their TCR under homeostasis, evidenced by Nur77^GFP^ expression and basal proliferation, yet are refractory to ex vivo restimulation. This uncoupling of TCR signaling from effector output distinguishes these cells from both exhausted and conventionally resting T cells. Strikingly, this homeostatic TCR activity is entirely IEC-dependent: loss of MHCIa on IECs nearly abolished TCR3^Rg^ pIEL differentiation and eliminated Nur77^GFP^ expression entirely. Cognate interactions with IECs are therefore required to establish the commensal-specific CD8αβ^+^ pIEL compartment. Thus, IEC pMHC presentation serves a dual purpose, driving differentiation and retaining commensal-specific CD8^+^ T cells within the epithelium, while simultaneously positioning them in proximity to epithelial signals that enforce restraint. How, then, do commensal-specific CD8αβ^+^ pIEL integrate opposing signals, continuous TCR engagement alongside potent coinhibitory and regulatory cues, to preserve tissue homeostasis while retaining functional potential? Part of the answer may lie in cell-intrinsic feedback: CD8αα and Nur77 are both upregulated downstream of TCR engagement and act to dampen further activation, providing a self-limiting brake that operates alongside coinhibitory receptors such as Lag3, Tim-3, and PD-1.^21,32,33,35,58–60,90,91^ Though some amount of IEC-directed cytotoxicity by commensal-specific pIEL may occur without overt pathology, the continuous presentation of microbial products likely necessitates the potent redundancy of restraint mechanisms observed here, where *Lag3* deletion alone is insufficient to break tolerance.^92,93^ Whether commensal-specific pIEL can be reactivated during infection or inflammation is an important open question; Type I interferons can lower TCR activation thresholds in intestinal T cells,^94^ raising the possibility that inflammatory conditions may license these cells to respond.

TGFβ signaling adds a further layer of restraint, potentially protecting pIEL from terminal effector differentiation.^73,74^ The low-level expression of stem-like markers (*Tcf7*, *Slamf6*, *Lef1*, *Id3*) in SFB-specific CD8αβ^+^ pIEL is consistent with this interpretation, though their functional significance remains to be determined. The requirement for both TCR and TGFβ signals to originate from the same cell is a critical feature of this model. Dendritic cell-mediated peripheral regulatory T cell induction requires coincident pMHCII presentation and αvβ8-dependent TGFβ activation by the same DC,^95,96^ a mechanism that ensures tolerogenic signals are delivered exclusively to cells receiving cognate antigen. Our findings extend this principle to the intestinal epithelium: IECs provide both the pMHCI signal that drives commensal-specific CD8^+^ T cell differentiation and αvβ6-activated TGFβ that enforces their functional restraint, ensuring that the cells most capable of epithelial cytotoxicity are precisely those most exposed to the signal that restrains them. That commensal colonization drives low-level local epithelial activation rather than systemic inflammation,^97,98^ likely also limits progression to exhaustion, preserving the long-term functional potential of these cells. The epithelial checkpoint identified here thus operates at multiple levels, shaping differentiation, enforcing restraint, and preserving long-term T cell fitness.

Prior studies have documented elevated CD8αβ^+^ T cell frequencies in the intestine of SFB-colonized mice;^4,16,99–101^ our work now identifies SFB as a major driver of antigen-dependent TCRαβ^+^CD8αβ^+^ pIEL expansion and establishes their antigen-specificity and epithelial regulation through development of a gut commensal-specific MHCIa-restricted TCR and tetramer. Although SFB has been largely characterized as a mouse commensal, recent studies reveal its presence in the human intestine.^102–104^ More broadly, we propose that the immunoregulatory mechanisms identified here are not SFB-specific but likely operate for any epithelium-adherent or mucus-penetrating member of the microbiota capable of eliciting continual presentation of pMHC complexes on IECs. As such, an important open question remains to identify other commensal microbes capable of eliciting CD8αβ^+^ pIEL responses. The clinical significance of this axis is underscored by the presence of anti-αvβ6 autoantibodies in UC,^82,83,87^ which would selectively disable the epithelial TGFβ checkpoint identified here.^85^ Loss of this restraint drives enhanced TCR sensitivity, cytokine production, and divergent inflammatory cell lineages in commensal-specific CD8^+^ T cells, transcriptionally aligned with IBD patient samples,^22^ suggesting that dysregulated pIEL responses to the microbiota may contribute to disease pathogenesis rather than simply accompany it. Together, these findings reframe intestinal tolerance not as an absence of commensal-specific immunity, but as an active, epithelially enforced restraint of commensal-specific CD8^+^ T cells and identify the αvβ6-TGFβ axis as a critical checkpoint whose disruption tips the balance from homeostasis to inflammation.

## Materials and Methods

### Ethics statement

All experiments were approved by the Institutional Animal Care and Use Committee of Benaroya Research Institute and performed according to their guidelines. All experimental work was performed with approval of the BRI Institutional Biosafety Committee.

### Mice

C57BL/6 (Jax 000664), CD45.1 Pep Boy (Jax 002014), CD45.1 JAXBoy (Jax 033076), B6 Vil1- Cre (Jax 004586), B6 Vil1-CreERT2 (Jax 020282), *Itgav-fl* (Jax 032297), *Rag2*^-/-^ *Il2rg*^-/-^ (Jax 014593), Nur77-GFP (Jax 16617), *Tcra*^-/-^ (Jax 002116), CD4-Cre (Jax 02217), Ai14 (TdTomato fate-mapper; Jax 007914), B6 Thy1^a^ (Jax 000406) mice were purchased from Jackson Laboratory. C57BL/6 mice were also purchased from Taconic Biosciences (Tac B6 B6-MPF) sourced from Germantown, NY as our SFB-colonized group. *H2-M3*^-/-^ mice were provided by Dr. Chyung-Ru Wang (University of Chicago, Chicago, IL).^45^ *H2-Kb transgene-fl* x MHCIa^-/-^ mice^57^ and H2-Kb^-/-^ H2-Db^-/-^ mice^43,48^ were generously provided by Dr. Aaron Johnson (Mayo Clinic Rochester, Minnesota) and Dr. Ellen Robey (University of California Berkeley, Berkeley, CA), respectively. All mice were housed at the Benaroya Research Institute in specific pathogen-free (SPF) conditions and were *Helicobacteraceae*-free as determined by PCR of fecal DNA (see below). Germ-free Swiss Webster mice were purchased internally from the University of Washington’s Gnotobiotic Animal Core (GNAC) and germ-free C57BL/6 mice were purchased from Charles River Laboratories; All germ-free mice and SFB-monoassociated mice were housed in the University of Washington Gnotobiotic Animal Core. Mice used in experiments were between 6 and 14 weeks of age at time of sacrifice unless specified. Both female and male mice were used for experiments and no sex differences were noted during analysis. Littermate controls were used to limit the variability and effects of microbiome. Mice were maintained at a temperature between 18°C and 23°C with 40–60% humidity and a 12 h-12 h light-dark cycle.

### In vivo treatments

#### Colonization

Segmented Filamentous Bacteria-containing fecal pellets from SFB-monoassociated gnotobiotic mice were a gift from Dr. Dan Littman (New York University Grossman School of Medicine, New York, NY). SFB-monoassociated feces were stored at -80°C until use. For colonization of germ-free mice or Jax mice, frozen fecal pellets were sterilely mashed and resuspended in sterile PBS and put through a 70µm cell strainer. The resultant slurry was used for oral gavage for fecal material transfer (FMT) – approximately 1 pellet/mouse was administered. To generate fecal donors for enriched-SFB (eSFB) flora, *Rag2*^-/-^ *Il2rg*^-/-^ (R2G2) mice were housed under SPF conditions and colonized using SFB-monoassociated fecal pellets as above. Fresh fecal pellets were collected from R2G2 mice and processed as above for de novo colonization with eSFB flora of SFB-free mice – for fresh feces, approximately 0.5 pellets/mouse was administered.

#### Tamoxifen administration

For activation of inducible Vil1-CreERT2, mice were given 4mg tamoxifen in corn oil by oral gavage 4 and 1 day prior to T cell transfer, then 2, 5, and 8 days post T cell transfer followed by euthanasia at day 14. Note: for SFB-specific T cell transfer experiments into *H2-Kb^Tg-flox^* x MHCIa^- /-^ mice, T cells were nucleofected with *H2-d1* Cas9-RNPs to avoid allograft rejection due to H2-Db expression on the transferred T cells (see nucleofection Method below). Knock down of H2- Db was confirmed by flow cytometry and knock down of H2-Kb on epithelial cells was also confirmed by flow cytometry on the CD45- fraction of the IEL samples.

For fate-mapping with CD103-CreERT2 and Ai14, mice were given 4mg tamoxifen in corn oil by oral gavage every other day starting at day 12 post T cell transfer and SFB colonization followed by euthanasia on day 21.

#### EdU administration and staining

To track proliferation and DNA synthesis, 0.5mg or 0.2mg EdU was injected i.p. in 200µL PBS. Mice were euthanized 24 hours later, and cell suspensions were prepared as described below. EdU staining was performed using the Click-iT EdU staining kit (ThermoFisher C10337 AF488), per manufacturer’s instructions with the following modifications/notes: permeabilization and fixation was performed using BD Cytofix/Cytoperm kit (BD 554655), Click-iT staining was performed in 100µL using half the amount of recommended Alexa Fluor Azide, and PE-based dyes were stained after Click-iT EdU staining (PE and PE-Cy7).

### ELISPOT

The mesenteric lymph nodes of Jax mice or from mice colonized with eSFB 10 days prior were harvested and mashed with 3% FBS complete RPMI through a 70µm cell strainer and CD8^+^ T cells were isolated using untouched EasySep™ Mouse CD8^+^ T Cell Isolation Kit (Stemcell Technologies 19853) per manufacturer’s instructions. CD11c^+^ antigen presenting cells (APCs) from spleens of SFB-free mice were isolated using mouse CD11c MicroBeads UltraPure (Miltenyi Biotec 130-125-835) after 25 min of enzymatic digestion with 100µg/mL Liberase TL (Sigma 5401020001) and 500µg/mL DNase I (Sigma DN25). 250,000 enriched CD8+ T cells and 250,000 enriched CD11c^+^ APCs were plated on a MultiScreen 96-well ELISPOT plate (Millipore MAIPS4510) that was coated with Mouse IFNγ ELISPOT Pair (BD 551881) per manufacturer’s instruction with 150µL/well in 10% complete RPMI. For treatments, anti-CD3 was used at 1µg/mL final concentration; one fecal pellet from SFB monoassociated gnotobiotic mice were resuspended in 200µL PBS and boiled for 1hr at 95°C and clarified at 100xg for 1 min and used at a final dilution of 1:250 or 1:500 in each well. Cells were incubated overnight before spots were developed per manufacturer’s instructions and read on an ImmunoSpot C.T.L. Analyzer and quantitated using ImmunoSpot.

### Quantification of bacterial DNA

Quantification of fecal DNA was performed as described.^40^ Briefly, fecal DNA was prepared using QIAGEN DNeasy PowerSoil Pro Kit (Qiagen 47016) according to the manufacturer’s instructions. For generating a standard curve of qPCR, SFB 16S rRNA gene fragment was cloned into the pCR™2.1-TOPO™ vector (ThermoFisher) and serial 10-fold dilutions from 10^9 copies/reaction of plasmid DNA was used. PCR containing primers (SFB736F 5′-GACGCTGAGGCATGAGAGCAT-3′ and SFB844R 5′- GACGGCACGGATTGTTATTCA-3′), 50ng of purified fecal DNA or plasmid standard, and PowerUp SYBR Green Master Mix in a 10μL reaction were mixed and the PCR was performed in a 7500 Fast Real-Time PCR System (Life Technologies). Fecal DNA from SPF mice was also screened for *Helicobacteraceae* by PCR (Pan *Helico* F: 5’-GCTATGACGGGTATCC-3’ and Pan *Helico* R: 5’-GATTTTACCCCTACACCA- 3’)^105^ and all mice used in the present study were negative.

### SABR epitope discovery

^51^TCR3 epitope discovery was enabled by using a signaling and antigen-presenting bifunctional receptors to encode MHC-I molecules presenting covalently linked peptides (SABR-I) for CD8^+^ T cell antigen discovery. H2-Kb SABRs were constructed using gene synthesis (Twist Biosciences) similar to the HLA-A0201 SABRs described previously. Retroviral vectors presenting LNGFR-2A-Stuffer-H2-Kb SABR were generated and used to clone epitope libraries. The H2-Kb SFB peptidome was predicted in silico by entering the SFB-Mouse-NYU (ncbitaxon:1073972) proteome into NetMHCI4.0 with predictions for 8-mer and 9-mer epitope lengths. The result was a pool of 456,594 8-mers and 455,129 9-mer peptides. This pool was further selected on peptides predicted to bind with a dissociation constant less than 500nM. The final pool was a library of 7,295 8-mer and 8,474 9-mer peptides covering 93% of the SFB proteome (theoretical max of predicted H2-Kb peptidome) as well as positive and negative control peptides. The libraries were encoded in single stranded oligonucleotide pools containing the following sequences (tggttacaggagggctcggca + reverse-translated-epitope + ^49–51–I–HF^ (NEB R3539) to remove the stuffer fragment. The oligonucleotide library was cloned using NEBuilder HiFi DNA Assembly (NEB E5520S) at over 30X coverage at the DNA level. Library quality control was performed by confirming the presence of epitopes in 24 colonies by Sanger sequencing (Azenta). The cloned library was used to transduce NFAT-GFP-Jurkat cells. NFAT-GFP, NFAT-GFP with SABR-SIINFEKL, and murine-CD8αβ-Jurkats and murineCD8αβ-Jurkats with OT-I TCR. Jurkats expressing TCR3 were made in-house. Retroviral production for Jurkat transduction and epitope discovery was performed as previously described.^63–65^ Briefly, VSV-G pseudo-typed retrovirus was produced by transfecting 293T with pHIT60, pMD2-G (VSV-G), and library transfer vectors using TransIT 293 (Mirus Bio MIR 2700) transfection reagent per manufacturers instructions. Jurkat cells were transduced as described using RetroNectin-coated plates (Takara T100A) and 4µg/ml protamine sulfate (MedChem^106^µm syringe filter and used directly without concentration, or frozen in -80^51,107(chap2)^ cells were co-cultured with TCR3 (in addition to control co-cultures) and humanCD69^+^GFP^+^ cells were sorted using a BD FACSAria Fusion. Genomic DNA was extracted as described using AMPure XP SPRI-beads (Beckman Coulter A63881) after cellular lysis with ChIP lysis buffer containing 1% SDS, 10mM EDTA in 50mM Tris-HCL pH8.1).^106^ After magnetic purification, 80% ethanol washes, and resuspension in H2O, total gDNA was used as template for serial PCR to amplify the integrated SABR DNA using KOD polymerase (Millipore KMM-101NV) as described.^51,107(chap2)^ The first round of PCR amplified the cloned epitope, and the second round of PCR added UDI-indexes and i5 and i7 adapters (see Table S2 for list of primers). PCR products were purified using AMPure XP SPRI-beads as above and DNA was quantified using a Qubit 4 Fluorometer and ran on a 2% agarose gel to ensure pure product amplification. DNA concentrations were adjusted appropriately and subjected to sequencing using the MiSeq Nano V2 kit (Illumina MS-103-1001) spiked with 25% PhiX DNA (Illumina FC-110-3001) per manufacturer’s instructions. Jurkat SABR library co-cultures were sequenced from two independent library transductions each with 3 replicate co-culture wells.

Enrichment scores for putative epitopes were calculated using linear regression models for relative epitope abundance in the cellular input and TCR3-selected humanCD69^+^NFAT-GFP^+^ samples. High confidence hits were selected based on their presence in all technical replicates and enrichment over input. The single SABR construct for SGLIYVTL was cloned from a gBlock purchased from Twist Biosciences using Gibson assembly.

### Retroviral production

Retrovirus for bone marrow and primary CD8^+^ T cell transduction was produced as described^108^ with some modifications – no chloroquine was used for transfection and virus was not concentrated. Briefly, mycoplasma-free 293T cells (ATCC CRL-3216) were cultured at 37°C, 5% CO_2_ in RPMI (Fisher cat. SH30027FS) supplemented with 100U/mL penicillin and streptomycin, 2mM glutamine, 10mM HEPES, and 55µM beta-mercaptoethanol with 10% FBS (Sigma 20C132) and plated at 60-70% density on Poly-L-Lysine-coated 15 cm tissue-culture-treated dish. The next day, transfection of pCL-Eco and the transfer vector was performed using 1:3 DNA:PEI (polyethylenimine) mixture. Culture media was changed 6 hours later, and virus was collected 48 hours from change, filtered through a 0.45µm syringe filter and used directly without concentration, or frozen in -80°C for later use. Retroviral transfer vectors for TCR overexpression were gifted to us by Dr. Shivani Srivastava (Fred Hutchinson Cancer Center, Seattle, WA) and are in the mp71 backbone modified to include a truncated Thy1.1 linked by P2A to track transduction by flow cytometry.

### Retroviral transduction of primary mouse CD8^+^ T cells and adoptive transfer

Primary mouse CD8^+^ T cell transduction was performed as described.^109^ Briefly, T cells from spleen and lymph nodes of B6 WT mice (Jackson Laboratories), or other strains as indicated, were isolated using untouched EasySep™ Mouse CD8^+^ T Cell Isolation Kit (Stemcell Technologies 19853) and cultured in 10% mTCM media (RPMI supplemented with 100U/mL penicillin and streptomycin, 2mM glutamine, 1mM sodium pyruvate, 10mM HEPES, and 55µM beta-mercaptoethanol with 10% FBS) with 50U/mL recombinant human IL-2 (Preprotech, AF-200-15) and activated using Mouse T-Activator anti-CD3/CD28 Dynabeads (Thermo Scientific 11452D) and cultured for 24-30 hours at 1 million cells/mL at 37°C (1:1 Dynabeads to T cells). Non-tissue culture treated dishes were coated with 12.5µg/mL Retronectin (Takara T100B) in PBS overnight at 4°C or at 37°C for 30 minutes. Retronectin-coated plates were washed with PBS then blocked with 2% BSA in PBS for 30 minutes at 37°C and coated with viral supernatants for 2 hours while being centrifuged at 3000xg at 32°C in a tabletop centrifuge the day of transduction. T cells and Dynabeads were harvested, counted, and cultured as above then centrifuged at 800xg at 32°C for 30 minutes and returned to the incubator. After 24 hours, T cells were harvested and resuspended in media containing 50 U/mL IL-2 as above. After another 24 hours, T cells were harvested and resuspended in mTCM media containing 10 U/mL recombinant human IL-15 (Preprotech, AF-200-15) and cultured for 48 hours. T cells were then removed from Dynabeads and rested 24 hours and transduction efficiency was determined by flow cytometry by staining for anti-Thy1.1 in the case of TCR overexpression. Between 50,000 and 500,000 transduced cells were adoptively transferred i.v. into recipient mice in PBS.

### Retrogenics/On Time

Retrogenic mice were generated as described with modifications^46^ using an “On Time” retroviral transfer vector gifted by Dr. Kristin Hogquist (University of Minnesota, Minneapolis, MN) modified to include P2A linked-expression of truncated Thy1.1 with TCRα under the control of CD4-Cre-mediated deletion of floxed GFP-stop^47^. In some cases, *Tcra*^-/-^ x CD4-Cre^+^ intercrossed with a Nur77-GFP transgene was used as donors to track TCR engagement. Briefly, bone marrow was harvested from *Tcra*^-/-^ x CD4-Cre^+^ donor mice, prepared as described^46^ and cultured in DMEM (Thermo Fisher Scientific 11995073) supplemented with 100U/mL penicillin and streptomycin,1X non-essential amino acids, 1mM sodium pyruvate, 10µM HEPES, and 55µM beta-mercaptoethanol with 20% FBS with 50ng/mL recombinant mouse SCF (Biolegend 579704), 50ng/mL recombinant mouse IL-6 (Biolegend 570804), 20ng/mL recombinant mouse IL-3 (Biolegend 575504) overnight at 37°C, 5% CO_2_ at about 3 million cells/mL in a petri dish. Non-tissue culture treated dishes were coated with 12.5µg/mL Retronectin (Takara T100B) in PBS overnight at 4°C, or at 37°C for 30 minutes. Retronectin-coated plates were washed with PBS then blocked with 2% BSA in PBS for 30 minutes at 37°C and coated with viral supernatants for 2 hours while being centrifuged at 3000xg at 32°C in a tabletop centrifuge the day of transduction. After 24 hours, cells were harvested, counted, and resuspended in media as above with about 6 million cells/well of a virus-coated plate at about 2 million cells/mL. Cells were then centrifuged at 800xg at 32°C for 30 minutes and returned to the incubator. After another 24 hours, cells were harvested, counted, and 3-5 million cells were injected i.v. into lethally irradiated Jax B6 recipients. A small portion of cells were cultured an additional day to determine transduction efficiency by flow cytometry using GFP as the transduction marker. Mice were bled 6-8 weeks later to determine engraftment of “On Time” transduced bone marrow.

### Bone Marrow Chimeras

Bone marrow was harvested from JAXBoy (CD45.1) mice as above, counted. Before injection, a T cell depletion was performed using anti-Thy1.2 (30-H12),^110^ anti-biotin microbead, and MACs columns. Cells were injected i.v., separately, into lethally irradiated H2-Kb^-/-^ H2-Db^-/-^ or Jax B6 recipient mice. For mixed bone marrow chimeras bone marrow was harvested from *Lag3*^-/-^(CD45.1, CD90.1) bones (a generous gift from Dr. E. John Wherry at the University of Pennsylvania, Philadelphia, PA, Jax: 026644) and B6 Thy1^a^ mice as congenic WT donors. Cells were counted, mixed 1:1 and injected i.v. into lethally irradiated Jax B6 recipients. In both cases, mice were rested for 8 weeks before use.

### Adoptive transfer of naive CD8^+^ T cells

Spleen and lymph nodes (except the mesenteric lymph node) of TCR3 On Time retrogenic mice were harvested and mashed through a 70µm cell strainer and CD8+ T cells were isolated using untouched EasySep™ Mouse CD8^+^ T Cell Isolation Kit (Stemcell Technologies 19853) with the addition of biotinylated anti-CD44 (clone IM7 used at 1:300) to enrich for naive CD8+ T cells (amount of selection cocktail and magnetic beads recommended by manufacturer were halved to enhance yield and purity). The frequency of Thy1.1^+^ cells was determined by flow cytometry and the volume was adjusted in PBS such that 10,000 SFB-specific CD8^+^ T cells were injected i.v. in 100µL in each mouse. On Time donors and recipient mice were CD45.2 unless stated in figure legends. Mice were then colonized with SFB, as described above, up to 24 hours later.

### Nucleofection of sgRNAs

Purified CD8^+^ On Time retrogenic cells were isolated as above or thawed from frozen stock and the frequency of Thy1.1^+^ cells was determined by flow cytometry and nucleofected using the P3 nucleofection kit as described.^35^ Predesigned sgRNA guides were purchased from IDT that target *Cd19* (5’-GGUGGAAUGCUUCAGACGUC-3’), *Itgae* (5’-ACAUUGGUCGGGAACCACAA-3’ and 5’-CAAGGGAGGCGUUAAUGUCC-3’), *Tgfbr2* (5’-ACGGCCACGCAGACUUCAUG- 3’ and 5’-GGACUUCUGGUUGUCGCAAG-3’), and *H2-d1* (5’-GGUGACUUCACCUUUAGAUC-3’ and 5’-UGUCGGCUAUGUGGACAACA-3’). Briefly, Cas9-RNP complex was made by combining 1-2µL of sgRNA (IDT) at 100µM and 6µg of Cas9 (ThermoFisher A36496) in 5µL of water and incubated at room temperature for 10min. Then 2-5 million cells were resuspended in P3 nucleofection reagent, Cas9-sgRNA RNP complex was added and cells were electroporated using Lonza 4D-Nucleofector system using DN-100 pulse code. Cells were incubated for 10min at 37°C in 5% CO_2_ with 100µL of warm completed RMPI, then washed, counted, and resuspended in PBS such that 50,000 CD8^+^ T cells were injected i.v. in 100µL in each mouse. Mice were then colonized with SFB, as described above, up to 24 hours later.

### Cell isolation

Cells from mesenteric lymph nodes and spleen were collected in 3% FBS complete media (RPMI supplemented with 100U/mL penicillin and streptomycin, 1X non-essential amino acids, 2mM glutamine, 1mM sodium pyruvate, 10mM HEPES, and 55µM beta-mercaptoethanol with 10% FBS) and mashed through a 70μm cell strainer to generate single cell suspensions (the spleen was subjected to ACK red blood cell lysis during mashing and then quenched with media). For intestinal preparations, where indicated in the text, the ileum was defined as the distal 10cm of the small intestine. Where segmentation was performed, the intestines were sectioned into thirds: duodenum, jejunum, and ileum, and large intestine including the cecal tissue. The lamina propria and IEL were prepared from each segment as follows: the intestine was dissected and collected in cold 3% FBS complete media Peyer’s patches and cecal patch was removed before the intestines were opened longitudinally, rinsed in PBS to clean intestinal contents off and incubation in 3% FBS complete media containing 5 mM EDTA (Sigma-Aldrich), 0.145 mg/ml DTT (Sigma-Aldrich) at 37°C for 20 minutes on a stirring platform. To separate the intestinal epithelial lymphocyte (IEL) and lamina propria (LP) layer, intestinal contents were transferred through a sterile fine-meshed kitchen strainer into a 500mL beaker. Strainer was tapped on beaker several times after straining and pieces of each intestine were transferred to a 50mL conical tube containing 10mL of 0% FBS complete media with 2mM EDTA. After shaking the tube vigorously for 30 seconds, the contents of the tube were strained again through the sieve into the beaker. The process was repeated three times. The tissue was minced finely in digest containing 0% FBS complete media with either 0.1mg/mL Liberase TL (Sigma 5401020001) and 0.5mg/mL DNase I (Sigma-Aldrich) or 1mg/mL Collagenase type VIII (Sigma C2139) and 0.25mg/mL DNase I, with continuous stirring at 37°C for 25 min. The fraction collected in the beaker was passed through a 100μm cell strainer and is the “IEL” fraction. Digested tissue for LP was passed through a 70μm cell strainer. Lymphocytes from each fraction were centrifuged at 550xg on a tabletop centrifuge and further enriched by centrifugation at room temperature at 695xg for 8 min after pellets were resuspended in 37.5% Percoll (GE Healthcare). After washing in 3% FBS complete RPI, cells were resuspended for downstream use.

### Ex vivo restimulation

Single cell suspensions from tissues were stimulated in 48-well, tissue culture-treated dishes in 300μL of 10% FBS complete media. For anti-CD3 restimulation, 1μg/mL final concentration was used and cells were incubated at 37°C, 5% CO_2_ for 1 hour before the addition of 1x GolgiPlug (BD BDB555029) and incubated an additional 3 hours. For phorbol myristate acetate (PMA) (Sigma P8139) and ionomycin (Sigma I0634) restimulation, 50ng/mL of PMA was used and 5μg/mL of ionomycin was used with the addition of 1x GolgiPlug and cells were incubated for 2.5 hours at 37°C, 5% CO_2_. After stimulation, cells were then washed with cold PBS and moved into a 96-well plate for antibody staining and analysis.

### Flow cytometry

Cells were resuspended in 96-well, U-bottom plates and stained in PBS containing Rat IgG, Fc Block, fixable live/dead (Thermo L23105) and cell surface antibodies (for list of antibodies, see Table S3) and incubated on ice for 25 minutes. Cells were washed with PBS and, where necessary, intracellular protein staining was performed using either a transcription factor staining buffer set (Thermo 00-5523-00) or BD Cytofix/Cytoperm kit (BD 554655). In most cases, counting beads (Polysciences 183285) were used to enumerate total cell counts. All cells were analysed by a BD LSRFortessa.

### Tetramers

TL tetramer staining to identify CD8αα-expressing cells was performed as described at a concentration of 5µg/ml for 30 minutes at 37°C in 3% FBS complete RMPI.^33^ Tetramer was procured from the NIH tetramer core. SGLIYVTL tetramer was made using the easYmer H2-Kb MHC tetramer kit (Eagle Biosciences 5004-01) and peptide purchased from Genscript. SGLIYVTL tetramer staining was performed with 10nM tetramer conjugated to PE-streptavidin (Agilent PJRS25-1) for 30 minutes at 37°C in 3% FBS complete RMPI. In both cases, tetramer staining was followed by a PBS wash and other cell surface markers.

### In vitro culture

Spleen and lymph nodes (except the mesenteric lymph node) of On Time retrogenic mice or TCR transgenic mice were harvested and mashed through a 70µm cell strainer and CD8+ T cells were isolated using untouched EasySep™ Mouse CD8^+^ T Cell Isolation Kit (Stemcell Technologies 19853) with the addition of biotinylated anti-CD44 (clone IM7 used at 1:300) to enrich for naive CD8^+^ T cells. CD11c^+^ antigen presenting cells (APCs) from spleens of *Tcra*^-/-^ or *H2-Kb^-/-^ H2-Db^- /-^* mice were isolated using mouse CD11c MicroBeads UltraPure (Miltenyi Biotec 130-125-835) after 25 min of enzymatic digestion with 100µg/mL Liberase TL (Sigma 5401020001) and 500µg/mL DNase I (Sigma DN25) in 0% FBS complete RPMI. Frozen or fresh 50,000 Thy1.1^+^CD8^+^ T cells and fresh 100,000 CD11c^+^ APCs were mixed in a 96-well U-bottom plate and cultured in 150µL of 10% FBS complete RPMI with various treatments added: anti-CD3 1µg/mL final concentration; all fecal pellets were resuspended in 200µL PBS and boiled for 1hr at 95C and clarified at 100xg for 1 min, and used at a final dilution of 1:250 or 1:500 in each well. SFB from cecal contents of SFB-monoassociated gnotobiotic mice were density-enriched and sorted as described,^38^ where 750,000 FACS events were resuspended in 300µL PBS and boiled for 1hr at 95C, final dilution was used at 1:50 in each well. Cells were incubated for 72 hours at 37°C then analyzed by flow cytometry for activation with antibodies against CD44, CD62L, CD25, and CD69. For peptide stimulation of SGLIYVTL (Genscript) and SIINFEKL (Genscript) were resuspended in DMSO at 10mg/ml and diluted to the indicated concentration in PBS co-cultures were set up as above and incubated overnight prior to analysis by flow cytometry. To determine MHC-restriction, DC2.4 cell lines were used as antigen presenting cells and co-cultures were performed as above in flat-bottom plates. WT and MHC-I KO cell lines were gifts from Dr. Laurent Coscoy (UC Berkeley).^48^

### Microscopy and histology

Staining Swiss rolls was done as previously described.^35^ Briefly, the small intestine was dissected then flushed with cold PBS, splayed open and rolled into a Swiss roll, then fixed in 4% paraformaldehyde (PFA) for 4hr then placed in 30% sucrose overnight at 4°C. The next day rolls were shaken gently in 50% OCT in PBS for >2hr and then embedded in OCT and frozen at -80°C. Eight-16 μm sections were cut on a cryotome, dried and then frozen at -20°C. Slides were then fixed in ice-cold acetone for 10 min, dried and then blocked in PBS with 2.5% goat serum and 2.5% Donkey serum at room temperature for 1hr. Sections were stained in primary antibodies overnight at room temperature (anti-Thy1.1-Alexa Fluor 647 (1:200), anti-Ep-CAM FITC (1:800)). Stained slides were washed in PBS then mounted in ProLong Diamond Antifade reagent (Invitrogen P36961). Images were acquired using an SP5 confocal microscope (Leica) or Echo Revolve. Histology was performed on intestinal clippings fixed in 10% normal buffered formalin before paraffin embedding, slicing, and staining with hematoxylin and eosin and imaged on a Leica DMi1 with iPhone 13 mini through eyepiece. Pathology was scored blind using a modified Marsh-Oberhuber classification.^111^

### RNA sequencing

#### 10x Single cell RNA sequencing

To sequence CD8αβ^+^ T cells from SFB colonized mice, two Jax B6 and two H2-M3^-/-^ mice were de novo colonized with enriched SFB fecal samples by oral gavage. Four weeks later, the IEL from the last 10cm of the small intestine was prepared and each mouse was stained with oligo-hashtagged antibodies and other surface markers. CD8^+^CD44^+^CD62L^-^ T cells from each mouse were FAC sorted using a BD FACSAria Fusion, counted, and pooled equally and 15,000 cells total were loaded onto one channel with a 10x Genomics Chromium Controller per manufacturer’s instructions. Gene expression, feature barcode, and TCR libraries were generated with Next GEM 5’ v2 reagents and sequenced on an Illumina NextSeq 2000. Base calls were performed using Illumina RTA on the instrument. Libraries were processed to FASTQ files, gene and feature barcode UMI counts, and assembled TCR sequences, using 10x Genomics Cell Ranger v.6.1.1. Reconstruction of full TCR sequences from 10x VDJ outputs was performed using Stitchr software^112^ and V(D)J calls from sequencing. The full TCR sequence from the ten most clonally expanded TCRs (genotype agnostic) were used for retroviral re-expression in vitro and subsequent in vivo adoptive transfer. TCR1 was from a Jax B6 mouse and TCR3 was from an *H2-M3^-/-^* mouse **(Table S1)**.

To sequence SFB-specific CD8^+^ T cells from various tissues, 10,000 naïve TCR3^Rg^ cells were transferred into 10 Jax mice and de novo colonized with enriched SFB fecal samples by oral gavage. 11 days later, spleen and MLN were enriched for Thy1.1-APC using MACS column enrichment and anti-APC microbeads (Miltenyi 130-090-855) and the LP and IEL from the last 10cm of the small intestine were prepared without enrichment. All 10 mice were pooled and each tissue was stained with oligo-hashtagged antibodies and other surface markers. CD8^+^Thy1.1^+^ T cells from each tissue were FAC sorted using a BD FACSAria Fusion, counted, and pooled equally and 14,000 cells total were loaded onto one channel with a 10x Genomics Chromium Controller per manufacturer’s instructions. Gene expression and feature barcode libraries were generated with Next GEM 5’ v3 reagents, and sequenced on an Illumina NextSeq 2000. Base calls were performed using BCL Convert v.4.2.7 on Illumina BaseSpace. Libraries were processed to FASTQ files and gene and feature barcode UMI counts using 10x Genomics Cell Ranger v.9.0.0.

To sequence SFB-specific CD8^+^ T cells in a setting with perturbed TGFβ signaling, 10,000 naïve TCR3 cells were transferred into three *Itgav*^fl/fl^ and three *Itgav*^fl/fl^ *Villin*^Cre/+^ mice and de novo colonized with enriched SFB fecal samples by oral gavage. 28 days later, IEL from the last 10cm of the small intestine was prepared and each mouse was stained with oligo-hashtagged antibodies and other surface markers. CD8+Thy1.1+ T cells from each mouse were FAC sorted using a BD FACSAria Fusion, counted, and pooled such that 1/3 of the cellular input was from *Itgav*^fl/fl^ mice and 2/3 was from *Itgav*^fl/fl^ *Villin*^Cre/+^ mice to account for enhanced T cell heterogeneity found in *Itgav*^fl/fl^ *Villin*^Cre/+^ mice. Target cell numbers were loaded onto one channel with a 10x Genomics Chromium Controller per manufacturer’s instructions. Gene expression and feature barcode libraries were generated with Next GEM 5’ v3 reagents, and sequenced on an Illumina NextSeq 2000. Base calls were performed using BCL Convert v.4.2.7 on Illumina BaseSpace. Libraries were processed to FASTQ files and gene and feature barcode UMI counts using 10x Genomics Cell Ranger v.9.0.0.

Quality control, demultiplexing, projection, clustering, and gene expression comparisons were performed using the Seurat toolkit v.5.4.0.^113^ Cells were filtered based on empirically determined thresholds for minimum and maximum gene and UMI counts and maximum percent of UMIs derived from mitochondrial, hemoglobin, and ribosomal protein genes. Cells were further filtered based on expression of genes characteristic of non-T cells, to remove contaminating B cells, monocytes, and intestinal epithelial cells. Cells were assigned to source sample based on empirically determine thresholds for hashtag counts; any cells positive for multiple hashtags were excluded. UMAP parameters were optimized using scDEED.^114^ Trajectory analysis was performed using Slingshot v.2.18.0.^61^

#### Bulk RNA sequencing

To sequence CD8^+^ T cells from *Itgav fl/fl* mice, 10,000 naïve TCR3 cells were transferred into four *Itgav*^fl/fl^ and four *Itgav*^fl/fl^ *Villin*^Cre/+^ mice and de novo colonized with enriched SFB fecal samples by oral gavage. 14 days later, IEL from the last 10cm of the small intestine was prepared and stained with surface markers. 400 CD8^+^Thy1.1^+^CD103^+^ T cells from *Itgav*^fl/fl^ mice and 400 of each CD8^+^Thy1.1^+^CD103^+^ and CD103^-^ T cells from *Itgav*^fl/fl^ *Villin*^Cre/+^ mice were FAC sorted directly into SMARTseq buffer using a BD FACSAria Fusion. Full-length cDNA was amplified using the SMART-Seq v4 Ultra Low Input RNA kit (Takara Bio, San Jose, CA). Sequencing libraries were constructed using the NexteraXT DNA Library prep kit (Illumina, San Diego, CA). Library concentrations were quantified using a Qubit fluorometer (Thermo Fisher, Waltham, MA), and libraries were pooled based on relative concentration to achieve equal depth, and sequenced on an Illumina NextSeq 2000. Base calling and FASTQ generation was performed on Illumina BaseSpace using BCL Convert v.4.2.7. Reads were trimmed and filtered using fastp v.0.24.0.^115^ Trimmed FASTQs were aligned to mouse genome assembly GRCm38.91, using STAR v.2.7.11b.^116^ Gene counts were calculated using htseq-count,^117^ and quality metrics were calculated using Picard v.3.1.0 (http://broadinstitute.github.io/picard/) and FastQC v.0.12.1.^118^

To sequence CD8^+^ T cells from *Lag3*^-/-^ mixed bone marrow chimeras, chimeras were de novo colonized with enriched SFB fecal samples by oral gavage. 28 days later, LP and IEL from the last 10cm of the small intestine was prepared and stained with surface markers. 400 CD8^+^CD44^+^C62L^-^Thy1.1^+^CD45.1 T cells from *Lag3*^-/-^ chimeras and 400 of each CD8^+^CD44^+^C62L^-^Thy1.1^+^CD45.2 T cells from WT control chimeras were FAC sorted directly into SMARTseq buffer using a BD FACSAria Fusion. Full-length cDNA was amplified using the SMART-Seq v4 Ultra Low Input RNA kit (Takara Bio, San Jose, CA). Sequencing libraries were constructed using the NexteraXT DNA Library prep kit (Illumina, San Diego, CA). Library concentrations were quantified using a Qubit fluorometer (Thermo Fisher, Waltham, MA), and libraries were pooled based on relative concentration to achieve equal depth and sequenced on an Illumina NextSeq 2000. Base calling and FASTQ generation was performed on Illumina BaseSpace using BCL Convert v.3.10.4. Reads were trimmed and filtered using Trimmomatic.^119^ Trimmed FASTQs were aligned to mouse genome assembly GRCm38.91, using STAR v.2.7.11a.^116^ Gene counts were calculated using htseq-count,^117^ and quality metrics were calculated using Picard v.3.1.0 (http://broadinstitute.github.io/picard/) and FastQC v.0.12.1.^118^

Libraries were filtered based on total read counts, percent of reads aligned to genome, and median CV of coverage. Read counts were normalized using the trimmed mean of m-values,^120^ and differential gene expression was calculated using the limma package.^121^ Gene set enrichment analysis was performed using GSEA as implemented in the GSEA_R package with MSigDB hallmark gene sets.^122,123^

### Statistics

Statistical analysis was performed in Prism (GraphPad Software, v10). Unpaired, two-tailed student’s t tests were applied to determine the differences between two individual groups. For multiple group comparisons ANOVA was used. For statistical details of specific experiments, please see respective figure legends.

### Data and materials availability

All data needed to evaluate the conclusions in the paper are present in the paper or the supplementary materials.

## Acknowledgements

We thank the Benaroya Research Institute (BRI) animal facility staff; Dr. Adam Wojno, Adin Pierce and Dr. Caroline Stefani (BRI Cell and Tissue Analysis Core); Drs. Vivian Gersuk, Kimberly O’Brien and Quynh-Anh Nguyen (BRI Genomics Core), Drs. Stephanie Osmond and Thomas Edwards (BRI Bioinformatics Core) and Samantha Kimmel for technical assistance and Dr. Jisun Paik (University of Washington) for assistance with gnotobiotic experiments. We thank Dr. Dan Littman for generous provision of monoassociated SFB feces; Dr. Kristin Hogquist and Dr. Shivani Srivastava for retroviral vectors; Dr. Ellen Robey for *H2-Kb*^-/-^*H2-Db*^-/-^ mice, Dr. Tessa Bergsbaken for CD103-fate-mapping mice, Dr. Laurent Coscoy for DC2.4 cell lines deficient for MHCIa molecules, Dr. E. John Wherry for *Lag3*^-/-^ bone marrow cells and Dr. Aaron J. Johnson for *H2-Kb^Tg-flox^* mice. TL-tetramer reagents were obtained from the NIH Tetramer Core Facility. We thank the M.J. Murdock Charitable Trust for support of flow cytometry, histology, imaging and genomics resources at BRI. We thank Drs. Meghan Koch, Jhimmy Talbot, Jessica Hamerman and Daniel Campbell for critical reading of the manuscript. We thank Drs. Jhimmy Talbot and Meghan Koch for constructive feedback on this project.

## Author contributions

Conceptualization: TMO, OJH

Formal analysis: TMO, MJD, ALB, OJH

Methodology: TMO, SV, VKKM, BC, ARG, AJJ, AVJ

Investigation: TMO, MJD

Funding acquisition: TMO, SV, ALH, OJH

Project administration: OJH

Writing – original draft: TMO, OJH

Writing – review & editing: TMO, OJH

