## Supplemental Figures for "Coincident epithelial signals restrain commensal-specific CD8αβ^+^ T cells in the intestine"

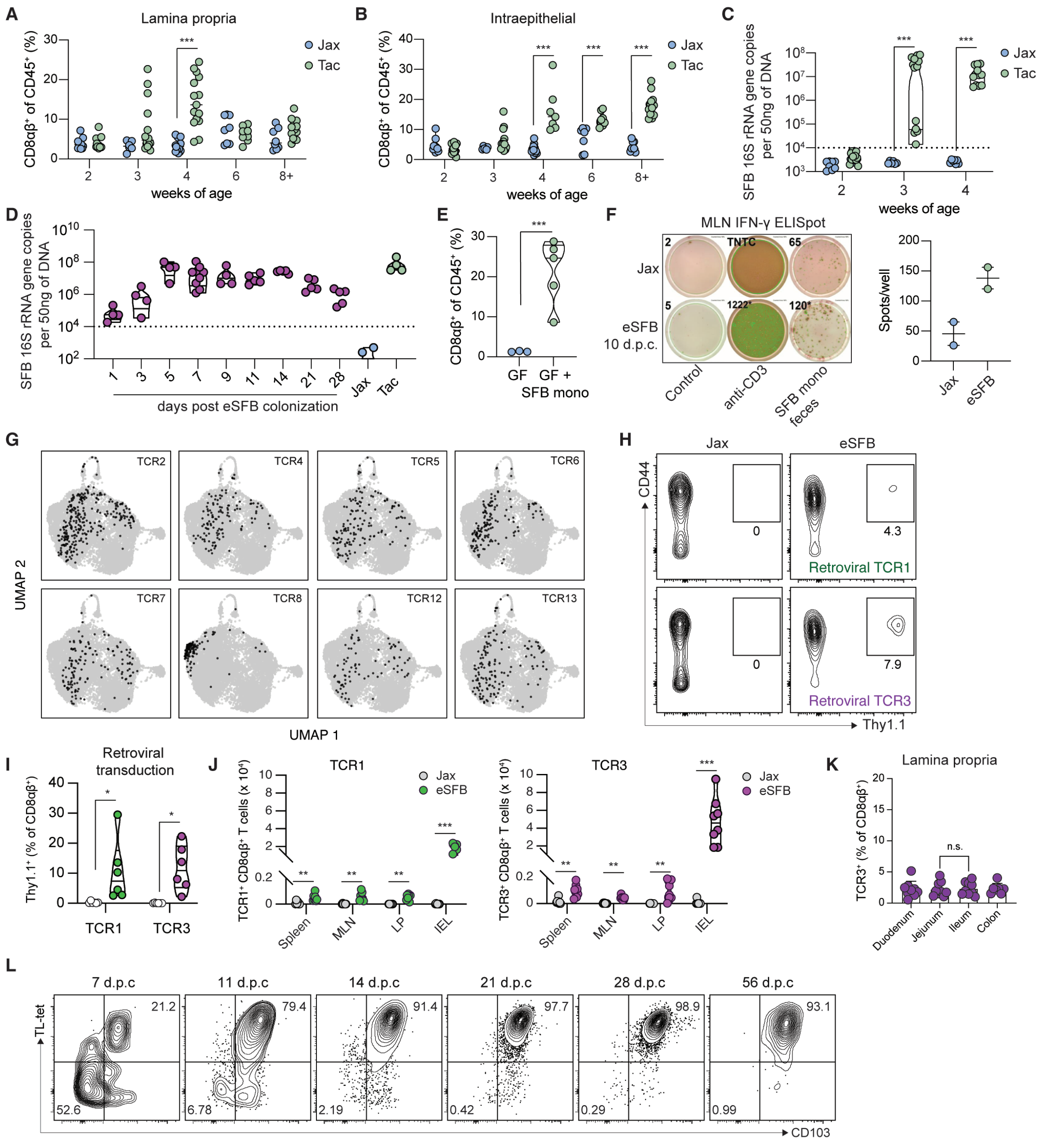
A
Lamina propria
CD8αβ+ of CD45+ (%)
weeks of age
Jax
Tac
B
Intraepithelial
C
SFB 16S rRNA gene copies per 50ng of DNA
D
days post eSFB colonization
E
GF
GF + SFB mono
F
MLN IFN-γ ELISpot
eSFB 10 d.p.c.
Control
anti-CD3
SFB mono feces
Spots/well
G
UMAP 1
UMAP 2
H
CD44
Thy1.1
Retroviral TCR1
Retroviral TCR3
I
Retroviral transduction
J
TCR1
TCR3
Spleen
MLN
LP
IEL
K
Lamina propria
Duodenum
Jejunum
Ileum
Colon
L
TL-tet
CD103

Supp 1.

**Fig. S1**. **(A)** Dot plot of CD8$\alpha\beta^+$ T cells as a percent of total live CD45$^+$ cells from the LP and **(B)** IEL fraction of the last 10cm (or, in the case of 2-week-old mice, the entire intestine) of the small intestine of mice originating from Jax and Tac at different age bred in-house. Data were collected from age matched mixed male and female Jax and Tac mice at each time point, n=6-14 per group per timepoint, pooled from five independent experiments. **(C)** Violin plot of qPCR analysis for total SFB load from feces from mice in (A). **(D)** Violin plots of qPCR analysis for total SFB load from feces over time from mice colonized with eSFB, n=4-5 per group. **(E)** Violin plot of CD8$\alpha\beta^+$ T cells as a percent of total live CD45$^+$ cells from germ-free Swiss Webster (SWR/J) mice colonized with SFB from monoassociated feces 28 days post colonization (d.p.c.), n=3-5 per group. **(F)** Purified CD8$^+$ T cells from the MLN of mice colonized with SFB for 10 days were incubated with purified CD11c$^+$ APCs with anti-CD3 or feces from SFB monoassociated mice overnight on an IFN-ɣ ELISPOT plate. One representative plate is shown with replicate dot plots, data are from two independent experiments. **(G)** Uniform manifold approximation and projection (UMAP) plot highlighting the cluster position and frequency of TCR clonotypes TCR2, TCR4, TCR5, TCR6, TCR7, TCR8, TCR12, and TCR13 from 10x single cell V(D)J sequencing of CD8$\alpha\beta^+$ pIEL from two Jax mice and two H2-M3$^{-/-}$ mice 28 d.p.c. with eSFB. **(H)** Representative flow plots of CD44 and Thy1.1 staining of total CD8$\alpha\beta^+$ T cells and **(I)** violin plots of CD44$^+$Thy1.1$^+$CD8$\alpha\beta^+$ pIEL as frequency of total CD8$\alpha\beta^+$ pIEL from mice adoptively transferred with 500,000 CD8$^+$ T cells transduced *ex vivo* with Thy1.1-TCR1 and Thy1.1-TCR3 retrovirus with or without eSFB 14 d.p.c. and T cell transfer, n=6 pooled from two independent experiments. **(J)** Violin plots of total retrogenic CD8$\alpha\beta^+$ T cells from spleen, MLN, LP and IEL of Jax mice receiving 10,000 naive TCR1$^{Rg}$ (green) and TCR3$^{Rg}$ (pink) CD8$^+$ T cells with or without eSFB 14 d.p.c and T cell transfer, n=7-8 per group from two independent experiments. **(K)** Bar graph of total TCR3$^{Rg}$ CD8$^+$ T cells from Jax mice receiving 10,000 naive cells 14 d.p.c. and T cell transfer. The small intestine was segmented in thirds and the colon was taken along with the cecum, n=9 mice per group from two independent experiments. **(L)** Representative flow plots of TCR3$^{Rg}$ CD8$^+$ T cells TL-tetramer (CD8αα) by CD103 from Jax mice receiving 10,000 naive cells with eSFB analyzed over time, n=8 per group from two independent experiments. P values were calculated using (A-D, H and J) ordinary one-way ANOVA with Tukey's multiple comparisons, (E) unpaired t test, and (K) quadrant frequencies plus/minus the standard deviation are shown. Statistical significance denoted as not significant (ns), * $P<0.05$, ** $P<0.01$, and *** $P<0.001$.

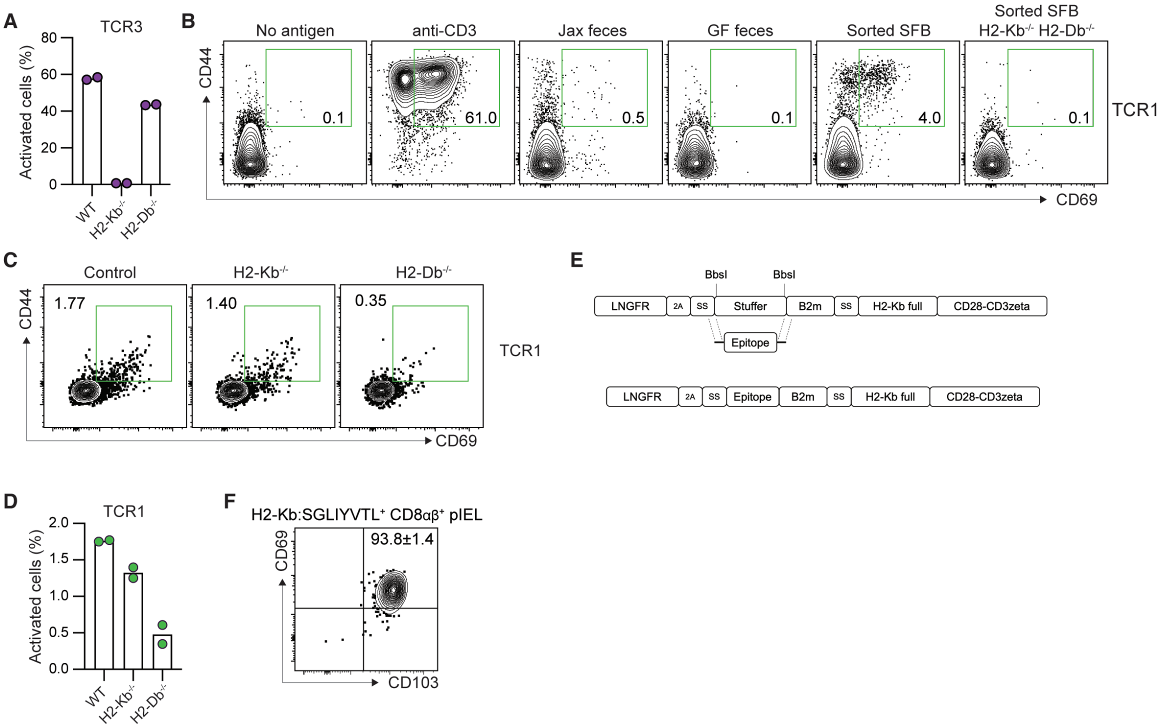
A
TCR3
Activated cells (%)
80
60
40
20
0
WT
H2-Kb-/-
H2-Db-/-
B
No antigen
anti-CD3
Jax feces
GF feces
Sorted SFB
Sorted SFB H2-Kb-/- H2-Db-/-
CD44
0.1
61.0
0.5
0.1
4.0
0.1
TCR1
CD69
C
Control
H2-Kb-/-
H2-Db-/-
1.77
1.40
0.35
TCR1
E
BbsI
BbsI
LNGFR
2A
SS
Stuffer
B2m
SS
H2-Kb full
CD28-CD3zeta
Epitope
LNGFR
2A
SS
Epitope
B2m
SS
H2-Kb full
CD28-CD3zeta
D
TCR1
Activated cells (%)
2.0
1.5
1.0
0.5
0.0
WT
H2-Kb-/-
H2-Db-/-
F
H2-Kb:SGLIYVTL+ CD8αβ+ pIEL
CD69
93.8±1.4
CD103

Supp 2.

**Fig. S2**. **(A)** Bar graphs from co-cultured naive TCR3$^{Rg}$ CD8$^{+}$ T cells and DC2.4 APC cell lines WT or deficient for individual MHCIa molecules treated with eSFB feces and incubated for 72 hours. Data are from two independent experiments. (**B)** Representative flow plots from co-cultured naive TCR1$^{Rg}$ CD8$^{+}$ T cells and purified CD11c$^{+}$ WT or MHCIa$^{-/-}$ antigen-presenting cells (APCs) treated with either anti-CD3, feces from Jax or germ-free mice, or FACS-purified SFB from monoassociated feces and incubated for 72 hours. Data were confirmed with two independent experiments. **(C)** Representative flow plots from co-cultured naive TCR1$^{Rg}$ CD8$^{+}$ T cells and DC2.4 APC cell lines WT or deficient for individual MHCIa molecules treated with eSFB feces and incubated for 72 hours. Data were confirmed with two independent experiments. **(D)** Bar graphs from co-cultured naive TCR1$^{Rg}$ CD8$^{+}$ T cells and DC2.4 APC cell lines WT or deficient for individual MHCIa molecules treated with eSFB feces and incubated for 72 hours. Data are from two independent experiments. **(E)** Schematic of Signaling and antigen-presenting bifunctional receptors (SABRs) epitope screening construct. **(F)** Representative flow plot of CD69 and CD103 staining on CD44$^{+}$SGLIYVTL-tetramer$^{+}$ CD8αβ$^{+}$ pIEL from Jax mice receiving 10,000 naive cells with eSFB, analyzed 14 d.p.c. and T cell transfer, n=7 per group pooled from two independent experiments. Quadrant frequencies plus/minus the standard deviation are shown.

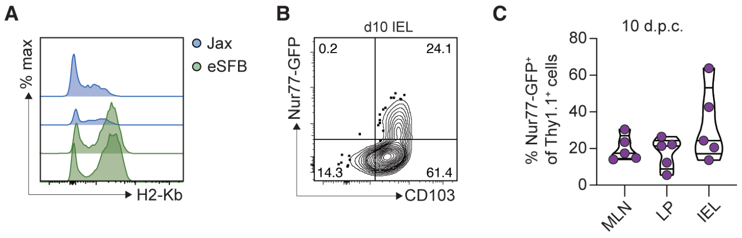
A
% max
Jax
eSFB
H2-Kb
B
d10 IEL
Nur77-GFP
0.2
24.1
14.3
61.4
CD103
C
10 d.p.c.
% Nur77-GFP+ of Thy1.1+ cells
80
60
40
20
0
MLN
LP
IEL

Supp 3.

**Fig. S3**. **(A)** Histograms of H2-Kb staining on live $CD45^{-}$ cells from the IEL fraction of the small intestine with or without eSFB 14 d.p.c. n=2. **(B)** Representative flow plots showing CD103 staining of $Nur77^{GFP}$ $TCR3^{Rg}$ $CD8^{+}$ T cells from ileum of eSFB colonized Jax mice 10 d.p.c. and T cell transfer, n=8-10 per group pooled from two independent experiments. **(C)** Violin plot of $Nur77^{GFP+}$ $TCR3^{Rg}$ $CD8^{+}$ T cells as a frequency of total $Nur77^{GFP}$ $TCR3^{Rg}$ $CD8^{+}$ T cells in mice from (B) at 10 d.p.c..

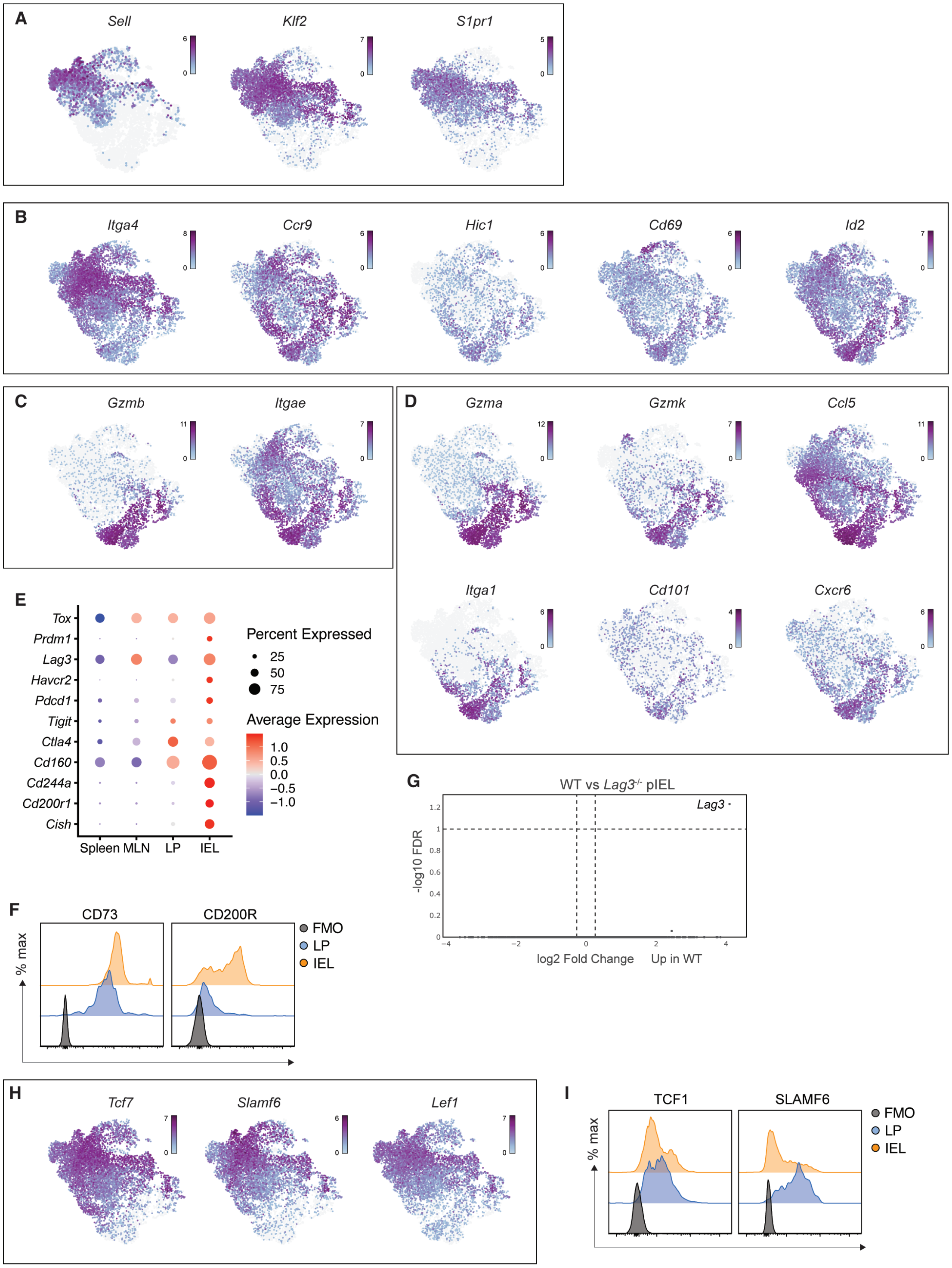
A
Sell
Klf2
S1pr1
B
Itga4
Ccr9
Hic1
Cd69
Id2
C
Gzmb
Itgae
D
Gzma
Gzmk
Ccl5
Itga1
Cd101
Cxcr6
E
Tox
Prdm1
Lag3
Havcr2
Pdcd1
Tigit
Ctla4
Cd160
Cd244a
Cd200r1
Cish
Spleen MLN LP IEL
Percent Expressed
25
50
75
Average Expression
1.0
0.5
0.0
−0.5
−1.0
F
CD73
CD200R
% max
FMO
LP
IEL
G
WT vs Lag3-/- pIEL
Lag3
-log10 FDR
log2 Fold Change
Up in WT
H
Tcf7
Slamf6
Lef1
I
TCF1
SLAMF6
% max
FMO
LP
IEL

Supp 4.

**Fig. S4**. **(A)** Uniform manifold approximation and projection (UMAP) plots showing log2 normalized expression of indicated secondary lymphoid organ homing genes, **(B)** gut homing genes, **(C)** pIEL associated genes, **(D)** effector genes, and **(E)** co-inhibitory receptor genes from 10x scRNAseq analysis of sorted $TCR3^{Rg}$ $CD8^{+}$ T cells from the spleen, MLN, ileum LP and IEL of Jax mice 11 d.p.c. and T cell transfer. Data were generated from cells pooled from 10 mice. **(F)** Protein expression of selected inhibitory receptors on $TCR3^{Rg}$ $CD8^{+}$ T cells from LP and IEL cells from Jax mice 14 d.p.c. and T cell transfer. **(G)** Volcano plot of differentially regulated genes from bulk RNA sequencing comparing sorted activated WT or *Lag3*$^{-/-}$ $CD8\alpha\beta^{+}$ pIEL from mixed bone marrow chimeras colonized with eSFB 28 days prior to analysis. **(H)** UMAP plots showing log2 normalized expression of indicated stem-like associated genes. **(I)** Protein expression of selected inhibitory receptors on $TCR3^{Rg}$ $CD8^{+}$ T cells from LP and IEL cells from Jax mice 14 d.p.c. and T cell transfer.

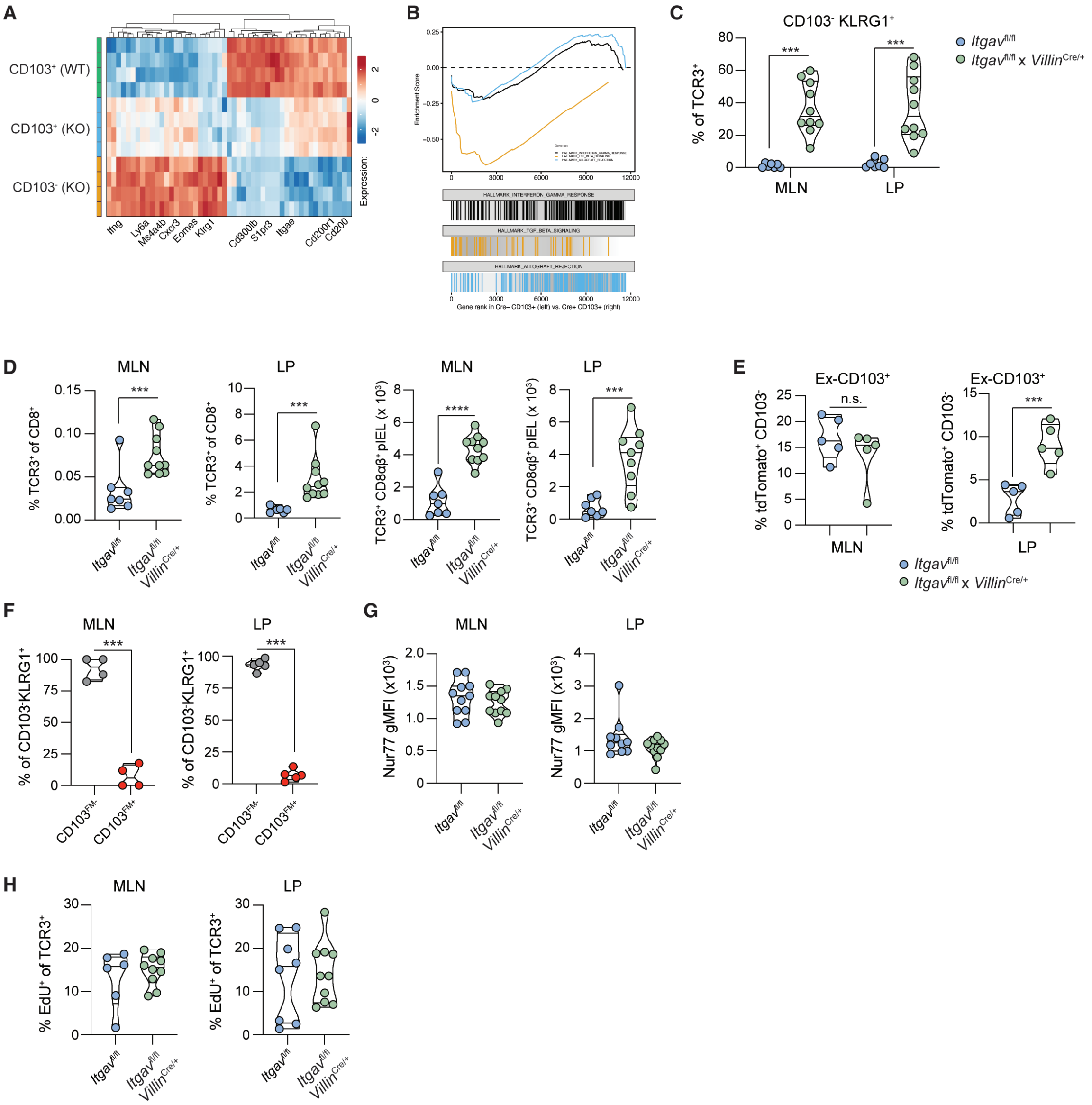
A
CD103+ (WT)
CD103+ (KO)
CD103- (KO)
Expression:
2
0
-2
Ifng
Ly6a
Ms4a4b
Cxcr3
Eomes
Klrg1
Cd300lb
S1pr3
Itgae
Cd200r1
Cd200
B
Enrichment Score
0.25
0.00
−0.25
−0.50
0
3000
6000
9000
12000
HALLMARK_INTERFERON_GAMMA_RESPONSE
HALLMARK_TGF_BETA_SIGNALING
HALLMARK_ALLOGRAFT_REJECTION
Gene rank in Cre− CD103+ (left) vs. Cre+ CD103+ (right)
C
CD103- KLRG1+
% of TCR3+
80
60
40
20
0
***
***
MLN
LP
Itgav fl/fl
Itgav fl/fl x Villin Cre/+
D
MLN
LP
MLN
LP
% TCR3+ of CD8+
0.15
0.10
0.05
0.00
***
% TCR3+ of CD8+
10
8
6
4
2
0
***
TCR3+ CD8αβ+ pIEL (x 10³)
8
6
4
2
0
****
TCR3+ CD8αβ+ pIEL (x 10³)
8
6
4
2
0
***
Itgav fl/fl
Itgav fl/fl Villin Cre/+
E
Ex-CD103+
Ex-CD103+
% tdTomato+ CD103-
25
20
15
10
5
0
n.s.
% tdTomato+ CD103-
15
10
5
0
***
MLN
LP
Itgav fl/fl
Itgav fl/fl x Villin Cre/+
F
MLN
LP
% of CD103-KLRG1+
100
75
50
25
0
***
% of CD103-KLRG1+
100
75
50
25
0
***
CD103 FM-
CD103 FM+
G
MLN
LP
Nur77 gMFI (x10³)
2.0
1.5
1.0
0.5
0
Nur77 gMFI (x10³)
4
3
2
1
0
Itgav fl/fl
Itgav fl/fl Villin Cre/+
H
MLN
LP
% EdU+ of TCR3+
30
20
10
0
% EdU+ of TCR3+
30
20
10
0
Itgav fl/fl
Itgav fl/fl Villin Cre/+

Supp 5.

**Fig. S5**. **(A)** Heat map of normalized expression of top 30 differentially up- and down- regulated genes between TCR3$^{Rg}$ CD8$^{+}$ T cells that are CD103$^{+}$ from *Itgav*$^{fl/fl}$mice and CD103$^{+}$ and CD103$^{-}$ from *Itgav*$^{fl/fl}$ *Villin*$^{Cre/+}$ littermate mice 14 d.p.c. and T cell transfer. **(B)** Gene set enrichment analysis (GSEA) of MSigDB hallmark gene sets indicated between CD103$^{+}$ from *Itgav*$^{fl/fl}$ and CD103$^{-}$ from *Itgav*$^{fl/fl}$ *Villin*$^{Cre/+}$ cells from (A). **(C)** Violin plots of CD103$^{-}$KLRG1$^{+}$ TCR3$^{Rg}$ CD8$^{+}$ T cells from MLN and LP as a frequency of total TCR3$^{Rg}$ pIEL from *Itgav*$^{fl/fl}$ *Villin*$^{Cre/+}$ or *Itgav*$^{fl/fl}$ littermate control mice 28 d.p.c. and T cell transfer, n=7-10 per group pooled from three independent experiments. **(D)** Violin plots of TCR3$^{Rg}$ CD8$^{+}$ T cells in MLN and LP plotted as total cells and as a frequency of total T cells from mice in (C). **(E)** Violin plots of CD103$^{-}$ tdTomato$^{+}$ cells from *Itgav*$^{fl/fl}$ *Villin*$^{Cre/+}$ or *Itgav*$^{fl/fl}$ littermate control mice in MLN and LP from CD103-CreERT2 fatemapping CD8$^{+}$ T cells transduced *ex vivo* with TCR3 prior to transfer into *Itgav*$^{fl/fl}$ *Villin*$^{Cre/+}$ or *Itgav*$^{fl/fl}$ littermate control mice. Mice were given 4mg tamoxifen every other day starting 12 d.p.c. and T cell transfer and analyzed day 21, n=5 per group. **(F)** Violin plot of frequency of CD103$^{-}$KLRG1$^{+}$ TCR3$^{Rg}$ CD8$^{+}$ T cells that are tdTomato$^{-}$ or tdTomato$^{+}$ from MLN and LP from *Itgav*$^{fl/fl}$ *Villin*$^{Cre/+}$ mice from (E). **(G)** Violin plot of the geometric mean fluorescence intensity (gMFI) of GFP from Nur77$^{GFP}$ TCR3$^{Rg}$ CD8$^{+}$ T cells from MLN and LP of *Itgav*$^{fl/fl}$ *Villin*$^{Cre/+}$ or *Itgav*$^{fl/fl}$ littermate control mice 28 d.p.c and T cells transfer, n=11-12 per group pooled from two independent experiments. **(H)** Violin plot of EdU$^{+}$ TCR3$^{Rg}$ CD8$^{+}$ T cells from MLN and LP as a frequency of total TCR3$^{Rg}$ CD8$^{+}$ T cells from *Itgav*$^{fl/fl}$ *Villin*$^{Cre/+}$ or *Itgav*$^{fl/fl}$ littermate control mice 28 d.p.c and T cells transfer, n=7-9 per group pooled from two independent experiments. P values were calculated using (C and E) two-way ANOVA with Šídák's multiple comparisons, (D, F-I) unpaired t test. Statistical significance denoted as not significant (ns), *** P <0.001, and **** P <0.0001.

| TCR_name | cloneIdTcrGraphClonalExpansion | nCellsInClonotype | nCellsWithSequence | chain | cdr3 | v_gene | v_gene_annotated | d_gene | j_gene | j_gene_annotated | c_gene |
|---|---|---|---|---|---|---|---|---|---|---|---|
| TCR1a | WT-Mouse-2_clone240 | 483 | 214 | TRA | CAMRGYSNTNKVVF | TRAV16*01 | TRAV16*06 | | TRAJ34 | TRAJ34*02 | TRAC |
| TCR2a | WT-Mouse-2_clone240 | 483 | 302 | TRA | CGASSSFSKLVF | TRAV5-4 | TRAV5-4*02 | | TRAJ50 | TRAJ50*01 | TRAC |
| TCR1b/2b | WT-Mouse-2_clone240 | 483 | 478 | TRB | CASSQTANSDYTF | TRBV12-2 | TRBV12-2*01 | | TRBJ1-2 | TRBJ1-2*01 | TRBC1 |
| TCR3a | H2-M3KO-Mouse-1_clone21 | 373 | 330 | TRA | CAASSGGSNYKLTF | TRAV14D-1 | TRAV14D-1*01 | | TRAJ53 | TRAJ53*01 | TRAC |
| TCR4a | H2-M3KO-Mouse-1_clone21 | 373 | 129 | TRA | CATDRGSGGSNYKLTF | TRAV8D-1 | TRAV8D-1*01 | | TRAJ53 | TRAJ53*01 | TRAC |
| TCR3b/4b | H2-M3KO-Mouse-1_clone21 | 373 | 352 | TRB | CASSQETGGYEQYF | TRBV5 | TRBV5*01 | TRBD1 | TRBJ2-7 | TRBJ2-7*01 | TRBC2 |
| TCR5a | H2-M3KO-Mouse-1_clone294 | 206 | 155 | TRA | CAMREYTNTGKLTF | TRAV16D-DV11 | TRAV16D-DV11*02 OR TRAV16*01 | | TRAJ27 | TRAJ27*01 | TRAC |
| TCR5b | H2-M3KO-Mouse-1_clone294 | 206 | 196 | TRB | CASSLGGNSDYTF | TRBV12-2 | TRBV12-2*01 | | TRBJ1-2 | TRBJ1-2*01 | TRBC1 |
| TCR6a | WT-Mouse-1_clone29 | 198 | 156 | TRA | CAASYGNEKITF | TRAV10 | TRAV10*01 | | TRAJ48 | TRAJ48*01 | TRAC |
| TCR6b | WT-Mouse-1_clone29 | 198 | 191 | TRB | CASSPGQGNSDYTF | TRBV29 | TRBV29*01 | TRBD1 | TRBJ1-2 | TRBJ1-2*01 | TRBC1 |
| TCR7a | WT-Mouse-1_clone248 | 185 | 142 | TRA | CAMRDYTNTGKLTF | TRAV16N | TRAV16N*01 | | TRAJ27 | TRAJ27*01 | TRAC |
| TCR7b | WT-Mouse-1_clone248 | 185 | 185 | TRB | CASSGTANSDYTF | TRBV12-2 | TRBV12-2*01 | | TRBJ1-2 | TRBJ1-2*01 | TRBC1 |
| TCR8a | H2-M3KO-Mouse-2_clone37 | 173 | 131 | TRA | CALDYNQGKLIF | TRAV6D-4 | TRAV6D-4*01 | | TRAJ23 | TRAJ23*01 | TRAC |
| TCR8b | H2-M3KO-Mouse-2_clone37 | 173 | 170 | TRB | CASSDWGEYEQYF | TRBV13-1 | TRBV13-1*02 | | TRBJ2-7 | TRBJ2-7*01 | TRBC2 |
| TCR12a | WT-Mouse-1_clone242 | 172 | 93 | TRA | CAMREYTNTGKLTF | TRAV16D-DV11 | TRAV16*01 OR TRAV16D-DV11*02 | | TRAJ27 | TRAJ27*01 | TRAC |
| TCR12b | WT-Mouse-1_clone242 | 172 | 163 | TRB | CASSLGGNSDYTF | TRBV12-2 | TRBV12-2*01 | | TRBJ1-2 | TRBJ1-2*01 | TRBC1 |
| TCR13a | WT-Mouse-1_clone333 | 172 | 154 | TRA | CALGSYTNTGKLTF | TRAV6N-7 | TRAV6N-7*01 OR TRAV6N-7*04 | | TRAJ27 | TRAJ27*01 | TRAC |
| TCR13b | WT-Mouse-1_clone333 | 172 | 166 | TRB | CASSRTGDYAEQFF | TRBV12-2 | TRBV12-2*01 | TRBD1 | TRBJ2-1 | TRBJ2-1*01 | TRBC2 |

**Note for some clonally expanded TCRs, a full length sequence was not able to be generated using Stitchr software due to missing IMGT annotation and were omitted from analysis**

### Table S1. CDR3 and V(D)J sequences of CD8αβ⁺ pIEL from mice colonized with eSFB

| Name | Sequence (5'-3')* | Purpose |
|---|---|---|
| UDI-ClassI-Fwd-Mix-1 | ACACTCTTTCCCTACACGACGCTCTTCCGATCTTGGCCTGCTTTGTTTGC | PCR1 |
| UDI-ClassI-Fwd-Mix-2 | ACACTCTTTCCCTACACGACGCTCTTCCGATCTGGCCTGCTTTGTTTGCC | PCR1 |
| UDI-ClassI-Fwd-Mix-3 | ACACTCTTTCCCTACACGACGCTCTTCCGATCTCCTGCTTTGTTTGCCGTG | PCR1 |
| UDI-ClassI-Rev-Mix-1 | GTGACTGGAGTTCAGACGTGTGCTCTTCCGATCTCCTCCACCACCGCTACCTC | PCR1 |
| UDI-ClassI-Rev-Mix-2 | GTGACTGGAGTTCAGACGTGTGCTCTTCCGATCCTTCCTCCACCACCGCTACC | PCR1 |
| UDI-ClassI-Rev-Mix-3 | GTGACTGGAGTTCAGACGTGTGCTCTTCCGATCGCTTCCTCCACCACCGCTAC | PCR1 |
| UDI-Index1 F | AATGATACGGCGACCACCGAGATCTACACagcgctagACACTCTTTCCCTACACGACGCTCTTCCGATCT | PCR2 index and adaptors |
| UDI-Index2 F | AATGATACGGCGACCACCGAGATCTACACgatatcgaACACTCTTTCCCTACACGACGCTCTTCCGATCT | PCR2 index and adaptors |
| UDI-Index3 F | AATGATACGGCGACCACCGAGATCTACACcgcagacgACACTCTTTCCCTACACGACGCTCTTCCGATCT | PCR2 index and adaptors |
| UDI-Index4 F | AATGATACGGCGACCACCGAGATCTACACtatgagtaACACTCTTTCCCTACACGACGCTCTTCCGATCT | PCR2 index and adaptors |
| UDI-Index5 F | AATGATACGGCGACCACCGAGATCTACACaggtgcgtACACTCTTTCCCTACACGACGCTCTTCCGATCT | PCR2 index and adaptors |
| UDI-Index1 R | CAAGCAGAAGACGGCATACGAGATaaccgcggGTGACTGGAGTTCAGACGTGTGCTCTTCCGATC | PCR2 index and adaptors |
| UDI-Index2 R | CAAGCAGAAGACGGCATACGAGATggttataaGTGACTGGAGTTCAGACGTGTGCTCTTCCGATC | PCR2 index and adaptors |
| UDI-Index3 R | CAAGCAGAAGACGGCATACGAGATccaagtccGTGACTGGAGTTCAGACGTGTGCTCTTCCGATC | PCR2 index and adaptors |
| UDI-Index4 R | CAAGCAGAAGACGGCATACGAGATttggacttGTGACTGGAGTTCAGACGTGTGCTCTTCCGATC | PCR2 index and adaptors |
| UDI-Index5 R | CAAGCAGAAGACGGCATACGAGATcagtggatGTGACTGGAGTTCAGACGTGTGCTCTTCCGATC | PCR2 index and adaptors |
| UDI-Index6 F | AATGATACGGCGACCACCGAGATCTACACgaacatacACACTCTTTCCCTACACGACGCTCTTCCGATCT | PCR2 index and adaptors |
| UDI-Index7 F | AATGATACGGCGACCACCGAGATCTACACacatagcgACACTCTTTCCCTACACGACGCTCTTCCGATCT | PCR2 index and adaptors |
| UDI-Index8 F | AATGATACGGCGACCACCGAGATCTACACgtgcgataACACTCTTTCCCTACACGACGCTCTTCCGATCT | PCR2 index and adaptors |
| UDI-Index9 F | AATGATACGGCGACCACCGAGATCTACACccaacagaACACTCTTTCCCTACACGACGCTCTTCCGATCT | PCR2 index and adaptors |
| UDI-Index10 F | AATGATACGGCGACCACCGAGATCTACACttggtgagACACTCTTTCCCTACACGACGCTCTTCCGATCT | PCR2 index and adaptors |
| UDI-Index6 R | CAAGCAGAAGACGGCATACGAGATtgacaagcGTGACTGGAGTTCAGACGTGTGCTCTTCCGATC | PCR2 index and adaptors |
| UDI-Index7 R | CAAGCAGAAGACGGCATACGAGATctagcttgGTGACTGGAGTTCAGACGTGTGCTCTTCCGATC | PCR2 index and adaptors |
| UDI-Index8 R | CAAGCAGAAGACGGCATACGAGATtcgatccaGTGACTGGAGTTCAGACGTGTGCTCTTCCGATC | PCR2 index and adaptors |
| UDI-Index9 R | CAAGCAGAAGACGGCATACGAGATcctgaactGTGACTGGAGTTCAGACGTGTGCTCTTCCGATC | PCR2 index and adaptors |
| UDI-Index10 R | CAAGCAGAAGACGGCATACGAGATGACCTGAAGTGACTGGAGTTCAGACGTGTGCTCTTCCGATC | PCR2 index and adaptors |

**Table S2. Oligo sequences used for SABR epitope discovery insertion amplification.**

| Flow Cytometry | | | | |
|---|---|---|---|---|
| **Target** | **Clone** | **Fluor/conjugation** | **Vendor** | **Catalog #** |
| b2-microglobulin | A16041A | PE | Biolegend | 154503 |
| CD103 | 2E7 | PE | Biolegend | 121406 |
| CD103 | 2E7 | BV510 | Biolegend | 121423 |
| CD200R (OX2R) | OX-110 | PE | Biolegend | 123907 |
| CD25 | PC61 | PE | BioLegend | 102007 |
| CD39 | Duha59 | PE-Cy7 | Biolegend | 143805 |
| CD4 | RM4-5 | BUV737 | BD Horizon | 612843 |
| CD4 | RM4-5 | PerCP/Cyanine5.5 | Biolegend | 100540 |
| CD44 | IM7 | biotin | Biolegend | 103003 |
| CD44 | IM7 | Alexa Fluor 700 | Thermo Scientific | 56-0441-82 |
| CD45 | 30-F11 | BUV395 | BD | 564279 |
| CD45 | 30-F11 | APC-eFluor 780 | Thermo Scientific | 47-0451-82 |
| CD45.1 | A20 | BUV395 | BD | 565212 |
| CD45.2 | 104 | BV510 | BioLegend | 109838 |
| CD45.2 | 104 | APC | BioLegend | 109814 |
| CD62L | MEL-14 | BV421 | BioLegend | 104436 |
| CD69 | H1.2F3 | eFluor 450 | Thermo | 48-0691-82 |
| CD69 | H1.2F3 | APC | Thermo Scientific | 17-0691-82 |
| CD73 | TY/11.8 | PE-Cy7 | Biolegend | 127223 |
| CD8a | 53-6.7 | Alexa Fluor 700 | Thermo Scientific | 56-0081-82 |
| CD8b | H35-17.2 | BV605 | BD | 740387 |
| CD8b | YTS156.7.7 | PE | BioLegend | 126608 |
| H2-Db | KH95 | PE | BioLegend | 111507 |
| H2-Kb | AF6-88.5 | FITC | BioLegend | 116505 |
| human CD69 | FN50 | PE-Cy7 | Biolegend | 310911 |
| IFNg | XMG1.2 | eFluor 450 | Thermo Scientific | 48-7311-82 |
| KLRG1 | MAFA | APC | Biolegend | 138412 |
| KLRG1 | MAFA | FITC | BioLegend | 138410 |
| Lag3 (CD223) | C9B7W | PE | BioLegend | 125208 |
| Ly108 (SLAMF6) | 330 AJ | APC | Biolegend | 134609 |
| NGFR (CD271) | ME20.4 | APC | Biolegend | 345108 |
| NGFR (CD271) | ME20.4 | PE | Biolegend | 345106 |
| PD-1 (CD279) | RMP1-30 | PE-Cy7 | BioLegend | 109110 |
| PD-1 (CD279) | J43 | APC | Thermo | 17-9985-82 |
| TCF1 | S33-966 | PE | BD | 564217 |
| TCRb | H57-597 | PerCP/Cyanine5.5 | Thermo Scientific | 45-5961-82 |
| Thy1.1 (CD90.1) | OX-7 | Alexa Fluor 700 | BioLegend | 202528 |
| Thy1.1 (CD90.1) | OX-7 | APC | BioLegend | 202526 |
| Thy1.2 (CD90.2) | 30-H12 | BV765 | Biolegend | 105331 |
| Tim-3 (CD366) | RMT3-23 | APC | Biolegend | 119705 |
| Tox | TXRX10 | eFluor 660 | ThermoFisher | 50-6502-80 |
| Live/Dead | | UV excited/V emissi | Thermo | L34962 |

| Other | | | | |
|---|---|---|---|---|
| **Target** | **Clone** | **Fluor/conjugation** | **Vendor** | **Catalog #** |
| CD3 (activation) | 17A2 | NA | ThermoFisher | 14-0032-85 |
| Thy1.2 (CD90.2) (depletion) | 30-H12 | biotin | Biolegend | 105304 |

| Imaging | | | | |
|---|---|---|---|---|
| **Target** | **Clone** | **Fluor/conjugation** | **Vendor** | **Catalog #** |
| Ep-CAM (CD326) | G8.8 | Alexa Fluor 488 | BioLegend | 118210 |
| Thy1.1 (CD90.1) | OX-7 | Alexa Fluor 647 | Biolegend | 202508 |

| Sequencing Hashatgs | | | | |
|---|---|---|---|---|
| **Target** | **Clone** | **Fluor/conjugation** | **Vendor** | **Catalog #** |
| TotalSeq™-C0301 anti-mouse Hashtag 1 Antibody | | oligo | Biolegend | 155861 |
| TotalSeq™-C0302 anti-mouse Hashtag 2 Antibody | | oligo | BioLegend | 155863 |
| TotalSeq™-C0303 anti-mouse Hashtag 3 Antibody | | oligo | Biolegend | 155865 |
| TotalSeq™-C0304 anti-mouse Hashtag 4 Antibody | | oligo | Biolegend | 155867 |
| TotalSeq™-C0309 anti-mouse Hashtag 9 Antibody | | oligo | Biolegend | 155877 |
| TotalSeq™-C0310 anti-mouse Hashtag 10 Antibody | | oligo | BioLegend | 155879 |
| TotalSeq™-C0313 anti-mouse Hashtag 13 Antibody | | oligo | Biolegend | 155885 |
| TotalSeq™-C0314 anti-mouse Hashtag 14 Antibody | | oligo | Biolegend | 155887 |
| TotalSeq™-C0315 anti-mouse Hashtag 15 Antibody | | oligo | Biolegend | 155889 |
| TotalSeq™-C0316 anti-mouse Hashtag 16 Antibody | | oligo | Biolegend | 155891 |
| TotalSeq™-C0317 anti-mouse Hashtag 17 Antibody | | oligo | Biolegend | 113931 |
| TotalSeq™-C0318 anti-mouse Hashtag 18 Antibody | | oligo | BioLegend | 113929 |

**Table S3. List of antibodies used throughout the paper.**

#### References:

Citations only found in methods are #105-123
